# Dissecting human embryonic patterning during organogenesis reveals a rostral midline progenitor population for mesoendoderm induction

**DOI:** 10.64898/2026.11.07.765055

**Authors:** Ying Yang, Hongan Ren, Mengyang Lu, Guanghui Tan, Yixuan Wang, Qingyu Cai, Junyao Jiang, Chenyu Guo, Yixin Li, Naixin Chen, Hong Chen, Xia Li, Fei Han, Niya Zhou, Xia Lan, Rong Li, Shigehito Yamada, Guojun Sheng, Weike Pei, Taihang Liu, Leqian Yu, Lizhong Liu

## Abstract

Understanding how human organogenesis is accomplished via intricate spatiotemporal patterning of organ primordia requires spatial atlases at cellular resolution. However, existing human embryo spatial atlases are built using sequencing-based platforms such as Stereo-seq, which have limited spatial resolution and blurred anatomical boundaries, leading to misclassification across adjacent cells. Here, we integrate in situ imaging-based transcriptomics (Xenium) with transcriptome-wide spatial profiling (Visium-HD) of Carnegie stages 9-12 embryos to delineate the patterning of core embryonic tissues at single-cell resolution, including the brain, spinal cord, somites, neural crest, notochord, heart, and gut tube. A new finding enabled by spatial single-cell resolution is the identification of rostral midline progenitors (RMPs), a cell population co-expressing SOX2, TBXT, and OTX2. Transcriptionally, RMPs are distinct from both neuromesodermal progenitors (NMPs) and prechordal mesoderm, and in vivo clonal barcoding in mice indicates that RMPs share a common developmental origin with rostral neural ectoderm. To interrogate their functional significance, we leveraged our spatial transcriptome data to recapitulate RMP-like cells from human pluripotent stem cells and showed that these cells can induce epiblast-like cells to acquire mesendodermal and endodermal fates, while themselves adopting an ectodermal fate consistent with their in vivo counterparts. This work provides a 3D, stage-resolved molecular cell atlas of tissue patterning in human embryo that serves as a reference for in vitro models.

## INTRODUCTION

Carnegie stage (CS) 9-12 Human embryos at age 19-30 days post-conception represent a crucial transition from gastrulation to the onset of organogenesis. The human embryo transforms from a trilaminar disc into one with a recognizable body plan, during which the foundational architecture for organogenesis is established. The coordinated emergence of organ rudiments is influenced by spatiotemporal signals from midline structures^1,2^. For example, the prechordal plate and notochord pattern the ventral forebrain and floor plate^3–6^. Mesodermal subtypes are specified mediolaterally around the midline, with their anteroposterior distributions reflecting a temporal maturation gradient^7–9^. In contrast, the endoderm regionalizes along the anteroposterior axis as the gut tube folds and closes ventrally^10,11^. Consequently, this developmental window marks a period of heightened vulnerability to developmental disruption that leads to major birth defects. Despite its significance, our current understanding of early human organogenesis and midline development relies largely on morphological characterization^12–14^ and animal models^15–17^.

Single-cell RNA sequencing (scRNA-seq) has generated an unparalleled reference for understanding human gastrulation^18^, but lacks spatial context. Spatial transcriptomics has enabled mapping of gene expression in their anatomical context^19–26^, yet existing atlases lack the single-cell resolution necessary to distinguish adjacent cell types and identify rare cell populations. Here, we present single-cell 3D molecular atlases of CS9, 11 and 12 human embryos. We identify a molecularly distinct progenitor cell population within the rostral (anterior) midline of the embryo that shares features of axial mesendoderm and neuromesodermal progenitors (NMPs), yet remains transcriptionally and functionally distinct from either lineage. We establish an in vitro model of those cells based on our spatial analysis of embryos to enable functional interrogation.

## RESULTS

### Spatial transcriptomics of CS9-12 human embryos

We analyzed three embryos obtained from elective terminations of pregnancy from healthy donors, staged as CS9 (late), CS11, and CS12 (**Fig. 1a**, and Methods)^10^. Morphological evaluation confirmed that all specimens were morphologically normal, exhibiting stage-specific developmental features. Specifically, the CS9 embryo had formed 3 somites. The CS11 embryo displayed an open cranial neuropore and a C-shaped heart tube. The CS12 embryo exhibited nearly closed cranial and caudal neuropores, prominent otic vesicles, and looped heart (**Fig. 1a, and Supplementary Fig. 1a**).

**Figure 1.**
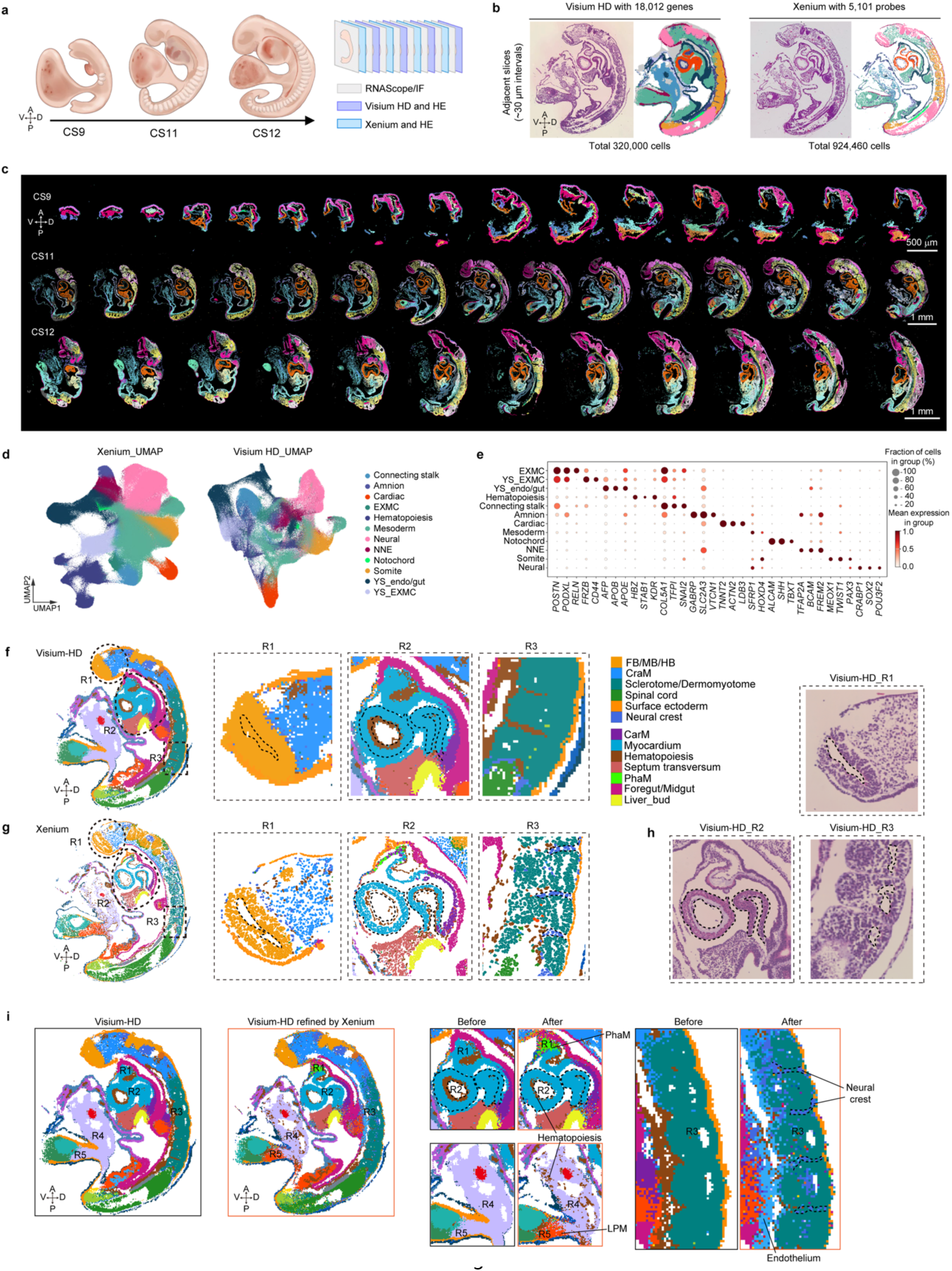
Spatial transcriptomics profiling of human embryos at CS9, CS11 and CS12 (a) Schematic overview of embryos at CS9, CS11, and CS12, illustrating key anatomical landmarks. Embryos were processed for spatial transcriptomics (Visium HD and Xenium), RNAscope, and immunofluorescence. (b) Representative Visium HD and Xenium sections from CS11 with corresponding hematoxylin and eosin (HE) stained sections. The total numbers of cells and genes detected by each platform are indicated. (c) Xenium sections from CS9, CS11, and CS12 with cell types color-coded. Extraembryonic tissues were detected but are excluded for clarity. Scale bars as indicated. (d) UMAPs of Visium HD and Xenium datasets of three stages. Cell types are color-coded. (e) Dot plot showing top marker genes for each identified cell cluster. Dot size represents the percentage of cells expressing each gene, and dot color indicates the average expression level (log-normalized counts) in expressing cells. (**f-g**) Spatial maps of Visium HD and Xenium. Full-section cell-type map with three regions (R1-R3, dashed outlines, indicated brain, heart, and somite, respectively) selected for magnification. Cell types are color-coded. (h) HE iamges of R1-R3 in Visium HD data showin in (f). (i) Representative iamges of Xenium-guided deconvolution of Visium HD data. Full-section cell-type maps before and after refinement (deconvolution) are shown. Heart, somite, and yolk sac are selected for magnification to present refinement detail. R1-R5 show representative regions of refinement. A, anterior; P, posterior; D, dorsal; V, ventral; AME, amniotic ectoderm; DE, definitive endoderm; EXM, extraembryonic mesoderm; Gas. Mes, gastro-intestinal mesentery; Inter. Meso, intermediate mesoderm; LB. Meso, limb bud mesoderm; LP. Meso, lateral plate mesoderm; Nas. Meso, nascent mesoderm; PSM, presomitic mesoderm; NE, neural ectoderm; NMP, neuromesodermal progenitor; NNE, non-neural ectoderm; Para. Meso, paraxial mesoderm; PGC, primordial germ cell; Pharyn. Endo, pharyngeal endoderm; Rostral Meso, rostral mesoderm; Splan.Meso, splanchnic mesoderm; VE, visceral endoderm; YE, yolk sac endoderm; YS. Meso, yolk sac mesoderm.

To profile single-cell gene expression within anatomical context, we performed Xenium in situ transcriptomics on 17 sagittal sections from the CS9 embryo, 19 from CS11, and 14 from CS12 (**Supplementary Table 1**). We used a 5,101-gene panel to target developmental regulators, signaling pathways, and core cellular processes (cell adhesion, proliferation, cytoskeleton, and metabolism etc.) (**Figs. 1b, c, Supplementary Table 2**). Cell segmentation was performed using a multimodal approach that combined DAPI-based nuclear detection with cell boundary markers (ATP1A1, CD45, CDH1) and interior signals (18S rRNA and αSMA/Vimentin)^27,28^. Deep learning-based algorithms were used to generate cell masks at single-cell resolution, integrating Z-stack information to resolve overlapping nuclei and ensure precise transcript assignment in densely packed tissues (**Supplementary Fig. 1c,** Methods). Spatial profiling was further validated by RNAscope and immunofluorescence (IF) staining on adjacent sections, which showed consistent expression patterns with the Xenium data (**Supplementary Fig. 1d**). To provide spatially continuous transcriptomic coverage and enable cross-validation of the Xenium data, in parallel we analyzed four sections adjacent to those used for Xenium using Visium-HD, a sequencing-based spatial transcriptomics platform that profiles the whole transcriptome across a tissue section at 2 μm resolution (**Fig. 1b, Supplementary Fig. 1b, and Supplementary Table 1**). The sections were subsequently subjected to histological staining. The resulting spatial maps preserved anatomical structures with high fidelity across developmental stages (**Figs. 1b-c**).

Through both platforms we detected 3.41×10^8^ transcripts and identified 1,244,460 cells across three embryos, after stringent quality control and filtering (Methods). Those cells were categorized into 12 major embryonic and extraembryonic lineages, including neural ectoderm (*SOX2⁺*, *POU3F2⁺*), non-neural ectoderm (NNE; *TFAP2A⁺*, *FREM2⁺*), notochord (*TBXT⁺*, *SHH⁺*), mesoderm (*TWIST1⁺*, *SFRP1⁺*), cardiac (*TNNT2⁺*, *ACTN2⁺*), somite (*MEOX1⁺*, *PAX3⁺*), amnion (*GABRP⁺*, *VTCN1⁺*), connecting stalk (*COL5A1⁺*, *TFPI⁺*), extraembryonic mesoderm (EXMC; *POSTN⁺*, *RELN⁺*), yolk sac endoderm/gut (YS_endo/gut; *AFP^+^*, *APOB^+^*), yolk sac extraembryonic mesoderm (YS_EXMC; *FRZB^+^*, *CD44^+^*) and hemogenic endothelial progenitors (HEP; *KDR⁺*, *HBZ⁺*) (**Figs. 1d-e, Supplementary Table 3**). Cell types identified by two platforms are concordant (**Supplementary Fig. 1e**). Subpopulation diversity expanded markedly from CS9 to CS12, suggesting this developmental window as a critical period for the onset of organogenesis (**Supplementary Fig. 1f**).

Comparison of matched anatomical structures in adjacent sections profiled by the two platforms revealed that Xenium provided sharper spatial segregation of neighboring cell types, whereas Visium-HD exhibited appreciable signal blending. For example, in the Visium-HD data, but not in the Xenium data, we detected the neural marker *SOX2* throughout the notochord, in a gradient spreading from the adjacent neural tube (**Supplementary Figs. 2a-d**), a pattern inconsistent with prior studies showing that the trunk level notochord is *Sox2*-negative^29,30^. Consistent with this limited boundary resolution, Visium-HD failed to faithfully resolve anatomical structures such as the rostral neuropore (R1), the atrial chamber (R2), and the somitocoel (R3) (**Fig. 1f**), all of which were resolved in fine detail by Xenium (**Fig. 1g**). Moreover, specific low-abundance cell types, such as endothelial cells within the atrial chamber (R2) and neural crest cells adjacent to somites (R3), were faithfully captured by Xenium but not by Visium-HD (**Figs. 1f-h, Supplementary Figs. 1g-h**).

Re-analysis of published Stereo-seq datasets from human^25^ and non-human primate embryos^31^ at comparable stages revealed similar boundary artifacts, including spurious SOX2 expression in the entire notochord (**Supplementary Figs. 2e-f**), and broadly dispersed cardiac and endothelial marker signals (**Supplementary Fig. 2g**). The original annotations of the published human Stereo-seq data primarily reflect large-scale tissue regions, with limited power to resolve local heterogeneity and low-abundance cell populations. In contrast, our Xenium data revealed these local, low-abundance cell types (**Supplementary Fig. 2h**). Together, these results indicate that signal blending, arising from lateral transcript diffusion and bin aggregation, is a general limitation of capture-based sequencing platforms such as Stereo-seq and Visium-HD^32,33^. In contrast, our Xenium dataset consistently yielded high-fidelity single-cell segmentation.

### Xenium-guided deconvolution enhances Visium-HD resolution and enables 3D atlas reconstruction

We leveraged the Xenium data to guide cell-type assignment in adjacent Visium-HD sections. We first deconvolved both datasets against human scRNA-seq reference^23,25^, using robust cell-type decomposition (RCTD)^34^. RCTD assigns cell-type identities to each spatial spot by estimating which reference cell types, and in what proportion, best explain that spot’s gene expression, while adjusting for platform-specific technical variance (**Supplementary Fig. 3a,** and **Methods**). This approach enabled finer resolution of local cell-type composition (**Supplementary Figs. 1g-h**). Extending this deconvolution approach to published CS12/CS13 Stereo-seq datasets^25^, also restored obscured cell populations, for example, endothelial cells in the heart region and neural crest cells adjacent to the somites (**Supplementary Figs. 3b-c**). Xenium most closely matched the cell-type composition measured by independent single-cell sequencing, with less ambiguity than Visium HD or the published Stereo-seq data (**Supplementary Figs. 3d-e, Supplementary Table 4**). Next, we performed further refinement of Visium-HD annotation using Xenium data only, employing Xenium cells directly as the cell-type reference to refine local cellular composition in adjacent Visium-HD sections. This Xenium data-guided refinement consistently improved the delineation of local cell-type boundaries and revealed lower-abundance cell populations that were not resolved in the original Visium HD annotations (**Fig. 1i, Supplementary Fig. 3f**).

Next, we sought to integrate cell-type identities and gene expression patterns with 3D tissue architecture. To this end, we computationally aligned serial Xenium sections using shared anatomical features (Methods)^35–37^, generating 3D maps for each stage (**Supplementary Fig. 4a, Movies 1-3**). These volumetric reconstructions encompassed germ-layer derivatives and extraembryonic tissues, with individual cells assigned unique spatial coordinates. The resulting 3D volumes are compatible with standard image processing software (e.g., ImageJ, Imaris), enabling 3D rendering of organ primordia morphology, assessment of progressive cell-type diversification within tissue architecture, and virtual re-sectioning along arbitrary planes (**Supplementary Fig. 4b).**

Together, we generated high-quality spatial transcriptomic atlases of human embryos spanning CS9-CS12 with single cell resolution. Leveraging this resource, we next investigated key development events shaping the human embryo.

### Spatial organization and pre-patterned HOX code of human neural crest

Neural crest cells are specified at the neural plate border, reside transiently in the dorsal neural tube, and then delaminate and migrate extensively to generate diverse derivatives, including much of the peripheral nervous system and the craniofacial skeleton, with distinct fates arising at different rostrocaudal (R-C) levels^38,39^. Whether this axial identity is pre-patterned within the dorsal neural tube before delamination or imposed by neighboring tissues during migration remains unresolved^40,41^, and direct evidence from early human embryos has been lacking.

Our data enable single-cell spatial mapping of neural crest populations relative to anatomical structures. UMAP analysis of neural crest cells from CS9, CS11, and CS12 embryos identified sub-populations: cycling cranial neural crest (cNC), migrating cNC, post-migratory (pharyngeal arch ectomesenchyme), pre-migratory neural crest (pNC), and Schwann cell precursors (SCPs) (**Fig. 2a, Supplementary Fig. 5a).** Mapping these populations onto 3D embryo maps revealed stereotyped positions. Specifically, cNC occupied the cranial region and pharyngeal arches, pre-migratory NC formed a continuous stripe along the dorsal neural tube, and SCPs localized along the trunk in the vicinity of somites (**Figs. 2b-c**) Along the R-C axis, pre-migratory NC cells already expressed a nested HOX code while residing in the dorsal neural tube, ordered from HOXA1/HOXA2 rostrally to HOXD10/HOXA13 caudally (**Fig. 2d**), indicating that rostrocaudal neural crest identity is pre-patterned before delamination in human embryos, rather than imposed solely by the migratory environment.

**Figure 2.**
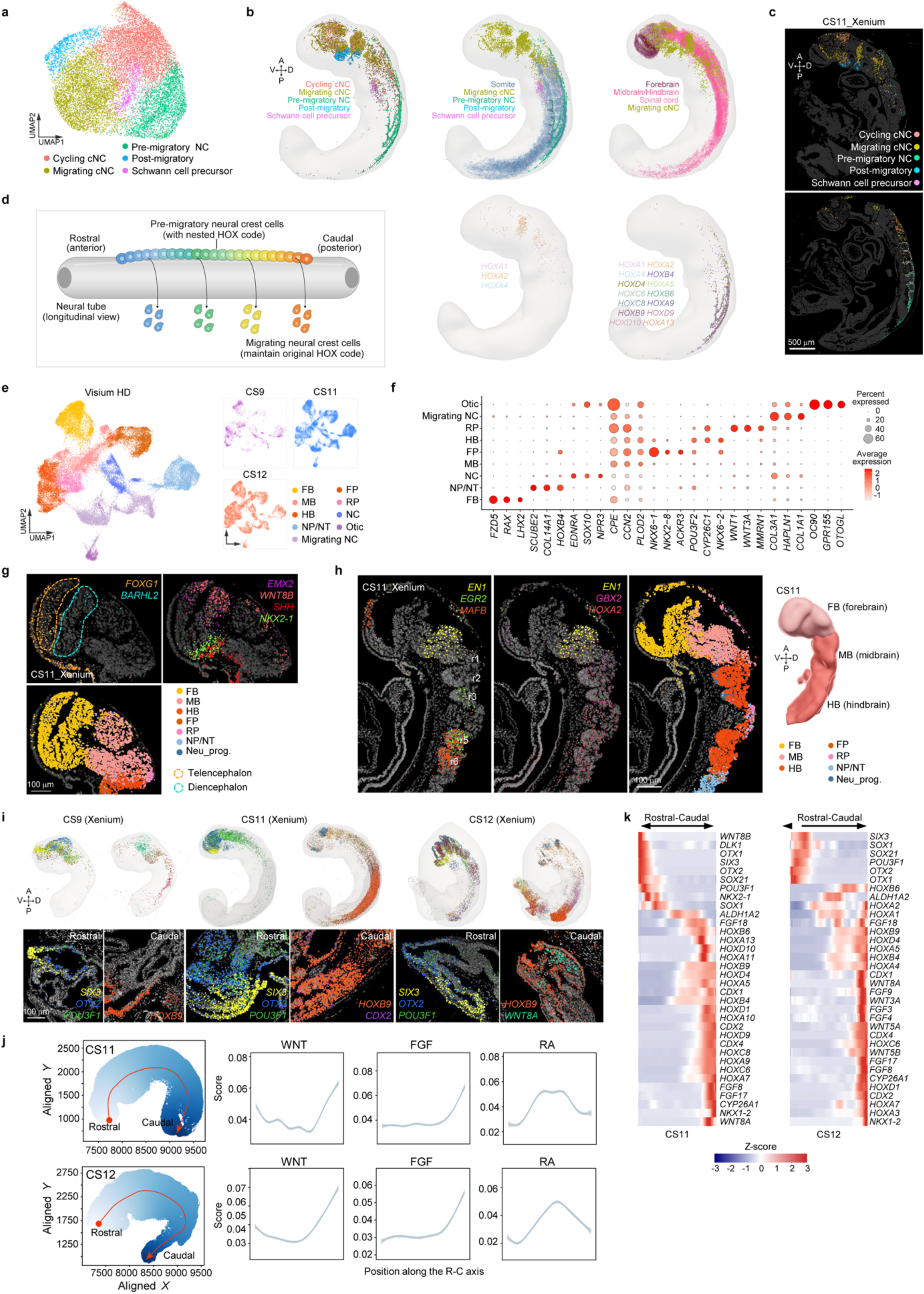
Neural crest patterning, neural tube regionalization, and primary brain vesicle formation (a) UMAP of neural crest cells of three stages, colored by cluster, including cycling/migrating craniofacial neural crest (cNC), pre-migratory neural crest (NC), post-migratory (pharyngeal arch ectomesenchyme), and Schwann cell precursor (SCP). (b) 3D embryo renderings: (upper left) all five populations; (upper middle) neural crest populations with somites; (upper right) migrating cNC relative to forebrain, midbrain/hindbrain, and spinal cord; (lower middle) HOX gene expression on migrating cNC; (lower right) nested HOX gene expression along the rostrocaudal axis in pre-migratory NC. (c) Xenium maps of neural crest subtype states on the CS11 embryo, showing different Z-planes. Scale bar, 500 µm. (d) Schematic shows the pre-patterning of neural crest cells. (e) UMAPs of ectodermal cell types, color-coded by cell type. Cells from CS9 (magenta), CS11 (blue), and CS12 (orange) are shown. (f) Dot plot showing marker gene expression across ectodermal cell types, derived from Visium HD sections. Dot size represents the percentage of cells expressing each gene; dot color indicates average expression level. (g) Spatial visualization of genes enriched in the forebrain at CS11, overlaid on Xenium sections. Cell types are color-coded. Orange and cyan dotted circles highlight the telencephalon and diencephalon, respectively. Scale bar, 100 µm. (h) Spatial visualization of genes enriched in the forebrain, midbrain, and hindbrain at CS11, overlaid on Xenium sections, with a corresponding 3D rendering of the CS11 brain vesicle. Rhombomeres (r) are indicated. Scale bar, 100 µm. (i) Expression of representative genes along the R-C axis at CS9, CS11, and CS12, displayed in 3D mesh models and corresponding Xenium sections. Scale bar, 100 µm. (j) Signaling pathway activities along the D-V and R-C axis at CS11 and CS12, quantified by integrated gene set pathway-associated transcriptional scores using irGSEA. (k) Heatmap of differentially expressed genes along the R-C axis at CS11 and CS12, ordered from rostral to caudal. NP, neural plate; NT, neural tube; NC, neural crest; FP, floor plate; RP, roof plate; FB, forebrain; MB, midbrain; HB, hindbrain; Otic, otic placode; Neu_prog., neuronal progenitor. A, anterior; P, posterior; D, dorsal; V, ventral.

### Neural tube regionalization and primary brain vesicle formation

The human central nervous system develops from the neural tube. Its patterning and closure occur concurrently at CS9-12^10,42^. Yet how positional identity is established along the emerging body axes during this window remains poorly understood. We therefore focused on the dynamics of cellular composition and signaling activities within the neural tube.

Firstly, our analyses recovered the cell populations expected from neural tube anatomy, including forebrain (FB, prosencephalon; *FZD5⁺*, *RAX⁺*), midbrain (MB, mesencephalon; *CPE⁺*, *CCN⁺*), hindbrain (HB, rhombencephalon; *POU3F2⁺*, *CYP26C1⁺*), neural plate/neural tube (NP/NT, *SCUBE2⁺*, *HOXB4⁺*), floor plate (FP, *NKX6-1⁺*, *NKX2-8⁺*), roof plate (RP, *WNT1⁺*, *WNT3A⁺*), neural crest (NC, *EDNRA^+^*, *SOX10^+^*), and migrating neural crest (*COL3A1⁺, HAPLN1⁺*) (**Figs. 2e-f**).

3D reconstruction placed above populations in their anatomical context, providing insight into the gene expression patterns underlying cell fate decisions and the morphogenesis (**Supplementary Figs. 5b-c**). At CS9, distinct expression domains were evident in the forebrain (*SIX3^+^FEZF1^+^OTX2^+^IRX1^low^*), midbrain (*SIX3^-^FEZF1^-^OTX2^+^IRX1^+^*), and hindbrain (*SIX3^-^ FEZF1^-^OTX2^-^IRX1*^+^). The midbrain-hindbrain boundary (MHB) was marked by *GBX2* expression, while *HOXA2* distinguished the hindbrain territory, which extended caudally to the first somite (**Supplementary Fig. 5f**). At CS11, the forebrain subdivided into telencephalic (*FOXG1⁺, EMX2⁺*) and diencephalic (*BARHL2⁺, OTX2⁺, POU3F1⁺*) domains (**Figs. 2g, i**), alongside the emergence of key signaling centers, including the mid-diencephalic organizer (MDO/ZLI), marked by *SHH* and positioned between the prospective prethalamus (*FEZF1/2⁺*) and thalamus (*IRX1⁺*)^43,44^, and the isthmic organizer (IsO) at the MHB^45^, marked by *FGF8* (**Supplementary Fig. 5g**). The hindbrain was segmented into rhombomeres (r). The metencephalon (r1) remained *HOX*-negative. *EGR2* marked r3/r5, and *MAFB* delineated the r5/r6 boundary (**Fig. 2h**). Whereas mouse *Hoxa2* occupies r2-r4 with an anterior boundary at the r1/r2 boundary and marks r4-derived neural crest entering the second arch^46,47^, HOXA2 in the CS11 human hindbrain was robustly detected in r5 and r6 on both platforms; this positional difference may reflect a stage difference, or a genuine species difference in rostral extent. By CS12, the midbrain had expanded substantially and the hindbrain had developed into eight rhombomeres (**Fig. 1c, Supplementary Fig. 5h**). The cranial neuropore remained incompletely closed at CS11. This coincided with broader expression of planar cell polarity (PCP) genes, including *CELSR1* and *FZD3/6*, than at CS9 (**Supplementary Figs. 5b-c**), consistent with a role of PCP signaling in driving neural fold elevation^48^. Technical limitations during tissue sectioning resulted in an incomplete forebrain of the CS12 embryo in this dataset (**Supplementary Fig. 5b**). Immunofluorescent staining of a stage-matched embryo confirmed the absence of the FOXG1^+^ forebrain region in the Xenium slide (**Supplementary Fig. 5d**).

We next investigated axial patterning within neural tube. Notably, the neural plate/tube at CS9 displayed a caudal-to-rostral gradient of maturation in its dorsoventral (D-V) patterning. Within the prospective brain, the anterior-most neural plate (*SIX3⁺*) remained *SHH*-negative, indicating that ventral patterning had not yet been initiated in the prospective anterior forebrain. By contrast, the *IRX1⁺IRX2⁺* posterior brain domain had initiated ventral patterning, with *SHH*, *NKX6-1*, and *FOXA2* all expressed in the ventral midline, consistent with floor-plate specification (**Supplementary Fig. 5e**). In contrast to the prospective brain, the prospective spinal cord had already established clear D-V patterning, with *PAX3* and *PAX7*confined to the dorsal domain (**Supplementary Fig. 5e**). At CS11, D-V patterning was well-established along the entire axis. *PAX3, PAX7, WNT3A,* and *WNT1* marked the dorsal neural tube. The floor plate was characterized by the expression of *NTN1*, a downstream target of the Sonic hedgehog (SHH) signaling pathway (**Supplementary Figs. 6a-c**). At this stage, opposing morphogenetic signaling were apparent, with high bone morphogenetic protein (BMP) signaling dorsally and elevated SHH signaling ventrally (**Supplementary Fig. 6d**). At CS12, the dorsal domain expressed high levels of *WNT3A, WNT8B,* and *EMX2*, whereas the ventral telencephalon was enriched for *NKX2-1*, marking the prospective ganglionic eminences. The dorsal midbrain exhibited expression of *WNT3A* and *WNT1* (**Supplementary Fig. 5h**). Neuronal differentiation was advanced in caudal regions, as evidenced by a contiguous zone of motor neuron precursors (*NKX6-1⁺, OLIG2⁺)* in the basal plate extending from the hindbrain to the cervical spinal cord (**Supplementary Figs. 6e-f**).

Regionalized gene expression along the R-C axis was detectable at CS9 and progressively refined by CS12 (**Fig. 2i**). The rostral-most domain was enriched for *SIX3, OTX1/2, SOX21*, and *POU3F1.* Along the trunk, HOX genes exhibited spatial collinearity: 3′ HOX genes (*HOXD4, HOXA4, HOXA5, HOXB4*) peaked more rostrally, and 5′ HOX genes (*HOXA9, HOXA13, HOXB9, HOXD9, HOXD10*) peaked progressively more caudally. The caudal-most end was defined by caudal growth zone signature genes, including FGF ligands, WNT ligands, *CDX2/4, NKX1-2*, the cell-cycle marker CDK1 (CDC2), and the RA-catabolizing enzyme *CYP26A1* (**Figs. 2i-k, Supplementary Figs. 5i-j**). Notably, the distribution of RA pathway components shifted along the R-C axis across stages. At CS9, CYP26A1 peaked in the rostral neural plate adjacent to the *SIX3⁺ OTX2⁺* rostral domain, while the RA-synthesizing enzyme *ALDH1A2* was enriched in the hindbrain and rostral trunk domain (*HOXA1⁺, HOXA2⁺, HOXB6⁺, FGF18⁺*) (**Supplementary Fig. 5j**). This arrangement is consistent with an RA-free zone that protects the prospective forebrain from posteriorization by trunk-derived RA^49^. By CS11-CS12, the CYP26A1 peak had relocated to the caudal end, while *ALDH1A2* remained enriched in the trunk (**Fig. 2k**). Consistent with this organization, pathway-activity scoring showed that RA-pathway pathway-associated transcriptional scores peaked in the trunk and were attenuated at both ends, whereas WNT and FGF scores peaked caudally (**Fig. 2j, Supplementary Fig. 5i**).

Together, our results show that positional identity in the human neural tube is established asynchronously along both body axes. Dorsoventral patterning matures in a caudal-to-rostral gradient, with the anterior forebrain patterned last, while rostrocaudal identity is refined through ordered brain domains, a collinear HOX code, and a caudal growth zone.

### Rostrocaudal patterning of the gut endoderm

We next investigated the rostrocaudal transcriptional program that governs endodermal patterning. Sub-regions of the primitive gut tube were already distinguishable at CS9, each defined by distinct marker genes (**Supplementary Figs. 6g-h**). Xenium spatial mapping showed that foregut, midgut, and hindgut domains were arranged rostral to caudal (**Supplementary Fig. 6i**). By CS11, newly emerged liver (*HABP2, RELN, SERPINA5, ALDH1A1, ONECUT3*) and lung (*NKX2-1, FOXE1, PAX8, ALCAM, FOXP2*) populations marked the onset of endodermal organ primordium specification. *NKX2-1⁺ FOXE1⁺* lung cells occupied the ventral foregut, *ALDH1A1⁺ HABP2⁺* liver cells occupied the hepatic region(**Supplementary Figs. 6h-i**). Of note, due to technical limitations only the CS11 sections captured the entire gut tube in this dataset. The midgut was lost in CS9 data, and hindgut was largely missing in CS12 data (**Supplementary Fig. 6g**).

### Spatiotemporal reorganization of cell-cell communication along the rostrocaudal axis

We further investigated cell-cell communication along the rostrocaudal axis by quantifying ligand-receptor interactions across tissue domains (**Supplementary Figs. 6j-k**). We observed a shift in signaling topology across stages. At CS9, cell-cell communication was regionalized to the anterior body parts between lateral plate mesoderm-surface ectoderm and the posterior body parts between the neural tube and surface ectoderm. By CS11, ectodermal and endodermal interactions were enriched anteriorly whereas mesodermal interactions were enriched posteriorly. By CS12, global signaling activity was enriched predominantly in the posterior region (**Supplementary Fig. 6k**). This spatiotemporal progression in cell-cell communications might reflect the rostrocaudal functional transition from the patterning of cranial structures to the secondary neurulation and paraxial mesoderm development in the caudal body parts^50–53^

### Axial patterning and morphogenesis of somites

Flanking the neural tube, the mesoderm is organized into paraxial and lateral plate mesoderm, which give rise to segmented somites and a chambered heart, respectively^10^. Yet how these adjacent mesodermal populations acquire distinct spatial identities and follow divergent morphogenetic trajectories in humans remains unclear.

We first focused on somite-related cell populations, including the presomitic mesoderm (PSM; *TBX6⁺, MSGN1⁺*), anterior PSM (A. PSM; *TBX6⁺, PCDH8⁺*), dermomyotome (*DMRT2⁺, PAX3⁺*), myotome (*MYOCD⁺, TRIM55⁺*), sclerotome (*FRZB⁺, SOX9⁺*), and mesenchyme (*TWIST2⁺, EBF2⁺*) (**Figs. 3a-b**). Spatial and pseudotime-trajectory analyses resolved continuous differentiation lineages from PSM progenitors to the sclerotome and dermomyotome/myotome compartments across CS9-CS12. (**Figs. 3a, 3c-f**), accompanied by stage-specific transcriptional programs (**Supplementary Figs. 7a-b**). Of note, because the caudal trunk becomes increasingly curved between CS9 and CS12, the assayed section series sample slightly different portions of the paraxial mesoderm at each stage. The apparent loss of A. PSM at CS12 reflects this sampling geometry rather than a biological loss.

**Figure 3.**
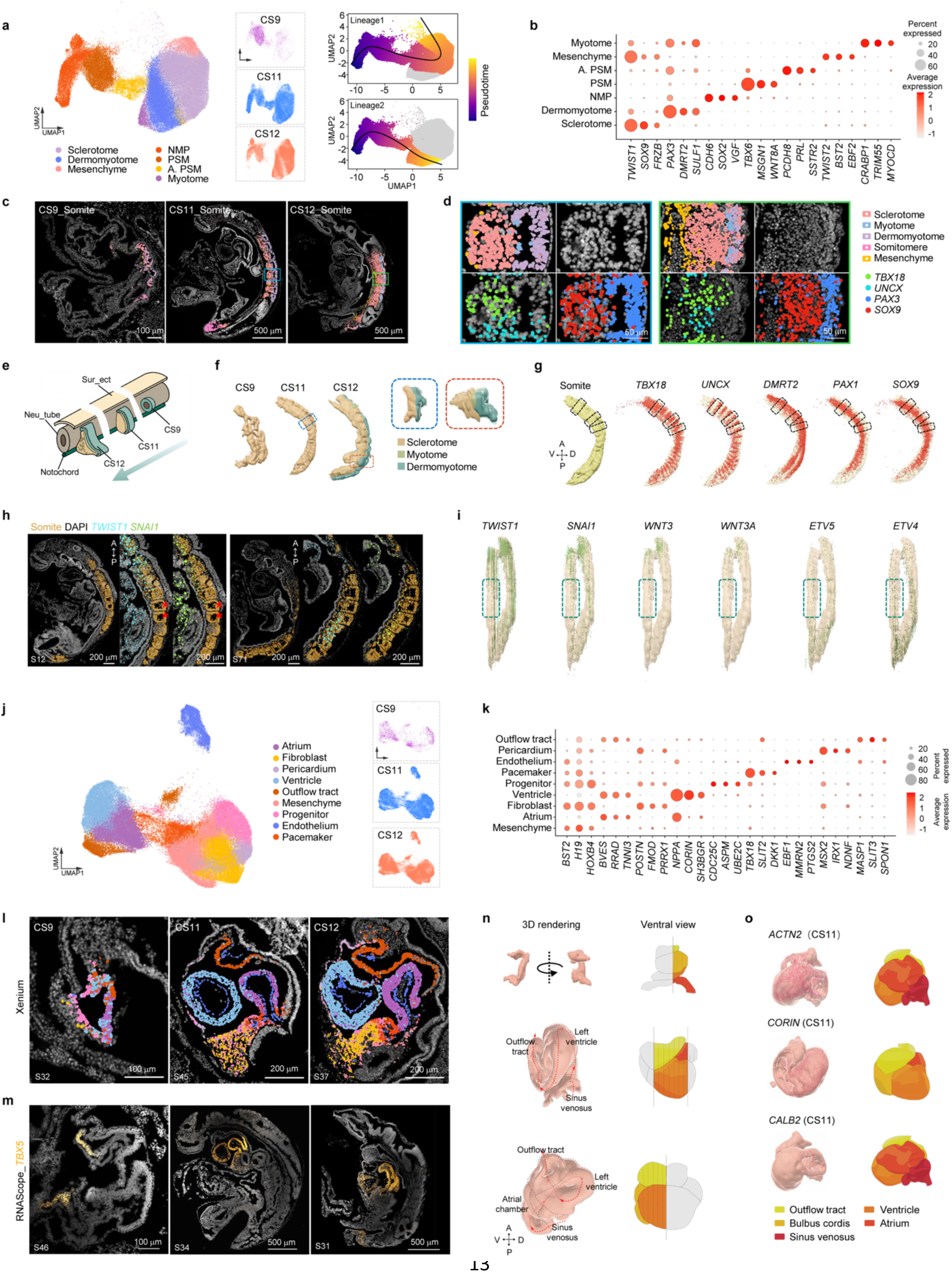
Patterning and morphogenesis of somite and heart (a) UMAPs of cell types associated with somitogenesis, color-coded by cell type. CS9 (magenta), CS11 (blue), and CS12 (orange) are shown separately on the right. The same embedding colored by inferred pseudotime (color scale) with the two reconstructed differentiation trajectories overlaid. both lineages originate in the NMP/PSM region and progress with increasing pseudotime toward the somite compartments. Lineage 1 toward sclerotome and Lineage 2 toward dermomyotome/myotome. **(b)** Dot plot showing marker gene expression across somitogenesis cell populations. Genes shown comprise the canonical markers used for annotation and the top differentially expressed genes of each population. The NMP population was annotated by co-expression of *TBXT* and *SOX2*. *CDH6* and *VGF* are shown as dataset-derived top differentially expressed genes that distinguish NMPs from the other somitogenesis populations; they are not claimed to be NMP-specific. Dot size represents the percentage of cells expressing each gene; dot color indicates average expression level. Cell types are shown on the y-axis; genes are ordered on the x-axis. (c) Representative Xenium sections showing somite-associated cell types at CS9, CS11, and CS12, color-coded by cell type. Scale bars as indicated. (d) Representative Xenium sections showing individual somites and expression of representative genes at CS11 and CS12. Scale bar, 50 µm. (e) Schematic summarizing somite development across CS9, CS11, and CS12. (f) 3D reconstructions of somites at CS9, CS11, and CS12, with cell types indicated. (g) 3D reconstruction of somites at CS11 with representative regionalized gene expression overlaid. Somitic cell types are displayed in 3D renderings for each stage, with representative genes marking cell identities along the dorsoventral and rostrocaudal axes. (h) Xenium sections at different Z-planes showing somites at occipital/cervical and thoracolumbar levels (left) and caudally positioned somites adjacent to the presomitic mesoderm (right). Two epithelial somites at thoracolumbar levels are highlighted by red asterisks. Magnified insets are shown to the right of each full section. Scale bar, 200 µm. (i) 3D reconstruction of somites at CS11, dorsal view. Expression of selected marker genes is shown. Epithelial somites are highlighted by dashed boxes. (j) UMAPs of cell types associated with cardiac development, color-coded by cell type. Cells are shown separately by developmental stage on the right. (k) Dot plot showing marker gene expression across cardiac cell populations. Dot size represents the percentage of cells expressing each gene; dot color indicates average expression level. (l) Representative Xenium sections from CS9, CS11, and CS12 showing cardiac cell types, color-coded cell types. Scale bars as indicated. (m) Representative RNAscope results showing *TBX5* expression at CS9, CS11, and CS12. Section numbers and scale bars as indicated. (**n**-**o**) 3D reconstructions of cardiac morphogenesis based on Xenium data, with representative genes highlighted. NMP, neuromesodermal progenitor; PSM, presomitic mesoderm; A. PSM, anterior presomitic mesoderm. A, anterior; P, posterior; D, dorsal; V, ventral.

At CS9, the PSM expressed the segmentation-clock gene *HES7*^54^, together with *MSGN1* and *TBX6*. Anterior to this domain, at the determination front within A. PSM, *MESP2* and *RIPPLY1* were sharply upregulated. Immediately anterior to A. PSM, three newly formed somites were marked by *MEOX1* and *TCF15*. Notably, Notch ligands *DLL1* and *DLL3* and the WNT readout *AXIN2* occupied spatially offset domains, consistent with the out-of-phase Notch and WNT oscillations that drive mesoderm segmentation^55–57^ (**Supplementary Fig. 7c**). Posterior to the determination front, the PSM enriched for *FGF8* and *WNT3A* and their downstream targets *DUSP6* and *SP5.* The anterior localization of *ALDH1A2* and the posterior restriction of *CYP26A1* likely established an anteroposterior retinoic acid (RA) gradient opposing the posterior FGF/WNT signaling center (**Fig. 2j**, **Supplementary Fig. 7d**).

By CS11 and CS12, somites underwent lineage segregation along rostrocaudal and dorsoventral axes. At CS11, the somitocoel was evident, and somites displayed anterior (*TBX18*⁺) and posterior (*UNCX*⁺) subdomains reflecting intrinsic somite polarity. The dorsolateral dermomyotome (*DMRT2*⁺, *PAX3*⁺) largely remained epithelial, enclosing the somitocoel, while the ventromedial sclerotome (*SOX9^+^, PAX1^+^*) was beginning epithelial-to-mesenchymal transition (EMT) (**Figs. 3d, g**). By CS12, the somitocoel had largely closed as sclerotome cells nearly completed EMT, generating *TWIST1/2⁺, FOXC1/2⁺, SOX9*⁺ mesenchyme (**Fig. 3b, d, Supplementary Figs. 7b, e, f**).

In avian embryos, sclerotomal differentiation initiates simultaneously across rostral (occipital and cervical) somites, whereas it proceeds in a rostral-to-caudal sequence in the trunk^58–61^. Whether comparable regional heterochrony exists in human embryos remains unknown. Unexpectedly, we found that trunk sclerotomal compartmentalization did not follow a strictly rostral-to-caudal progression in humans. Specifically, at CS11, whereas two to three thoracolumbar trunk somites retained epithelial morphology, more caudally positioned somites showed elevated *SNAI1* and *TWIST1* in their ventromedial domain (**Fig. 3h**). This indicates that, at thoracolumbar levels, younger caudal somites undergo EMT ahead of their older (more rostral) counterparts, suggesting that sclerotome differentiation tempo is uncoupled from the rostrocaudal sequence of somitogenesis. We further leveraged our 3D map to compare transcriptomic profiles across thoracolumbar somites (Somite-M), caudal somites (Somite-P), and rostral occipital/cervical somites (Somite-A). Thoracolumbar somites expressed high levels of epithelial junction-associated genes *CXADR* and *ALCAM*, consistent with their epithelial morphology, as well as *PAX3* and *ALDH1A2*, indicating dermomyotome identity and an RA-rich environment that maintains the epithelial somite state^62^ (**Fig. 2j, Supplementary Fig. 7g**). We then examined the WNT and FGF pathway activities, which are associated with EMT and somite maturation^63,64^. Notably, *WNT3* and *WNT3A* expression was low in these epithelial thoracolumbar somites, and FGF pathway activity, reflected by *ETV4* and *ETV5*, was also reduced relative to caudal somites (**Fig. 3i**). These findings suggest that regional heterochrony in somite development reflects position-dependent RA, WNT, and FGF signaling activities. Collectively, these results reveal that human trunk somite maturation is regionally heterogeneous, with tempo influenced by the local signaling environment rather than solely by somite birth order.

### Patterning and morphogenesis of cardiovascular structures

The heart progressed from a straight tube at CS9 to a C-shaped loop at CS11 and N-shaped structure by CS12 (**Supplementary Fig. 8a**). We identified cardiac progenitor (*NKX2-5^+^*, *ASPM^+^*, UBE2C⁺), ventricular myocardium (*CORIN*^+^, *NPPA*^+^), pericardium (*MSX2*^+^, *IRX1*^+^, *NDNF*^+^), endothelium (*MMRN2*^+^, *PTGS2*^+^) (**Figs. 3j-k**). At CS9, endothelial cells were not detectable. By CS11-CS12, endothelial cells had emerged progressively, and the heart had regionalized into chambers (**Figs. 3j, l**). Ventricular cells were enriched at the outer curvature and atrial cells (*TBX5*^+^) at the inflow region (**Figs. 3l, m**). Chamber-specific genes (*CORIN* and *CALB2*) were established, with a transitional zone co-expressing both atrial and ventricular markers at the atrioventricular canal (**Figs. 3l, 3o, and Supplementary Figs. 8a-b**).

The sinus venosus, comprising endocardial endothelium, mesenchymal cells, and pacemaker cells, was identifiable in the heart tube and incorporated into the developing atrium (**Figs. 3l, 3n-o**). At CS11, pacemaker cells resided adjacent to fibroblast and atrial myocardium (**Fig. 3l**), consistent with coordinated cardiac contraction by this stage^10,65^, and their co-localization suggests a common progenitor origin^66^. By CS11-CS12, endothelial networks became organized, with aortic sprouting extending dorsally to form segmental arteries, and endothelial cells appearing within the brain and neural tube (**Supplementary Fig. 8c**).

Using CellPhoneDB to infer candidate ligand-receptor interactions from spatially resolved expression (**Supplementary Fig. 8d**), we found strong myocardium-foregut endoderm signaling at CS9, mediated by *BMP2/4* and *FGF1/2/4*^67,68^, as well as interactions between cardiac lineages and the LPM, notochord, and neural crest (**Supplementary Figs. 8d-f**). At CS11-CS12, cardiac interactions with LPM, gut endoderm, and endothelium persisted. Notably, *WNT5A* and its receptor *FZD4* were expressed in the foregut endoderm and adjacent myocardium, respectively (**Supplementary Fig. 8f**), implicating non-canonical WNT signaling in outflow tract septation. Together, these findings suggest a role for myocardium-endoderm interactions in heart morphogenesis.

### Identification of rostral midline progenitors

UMAP analysis of the notochord niche revealed stage-distinct clustering of cells across CS9, CS11 and CS12 (**Fig. 4a**). While canonical notochord markers such as *TBXT*, *COL2A1*, *NOG*, and *SHH* displayed uniform expression along the notochord, the expression of a subset of genes showed marked rostrocaudal regionalization, suggesting distinct signaling environments in the rostral and caudal notochord domains (**Supplementary Fig. 9a**).

**Figure 4.**
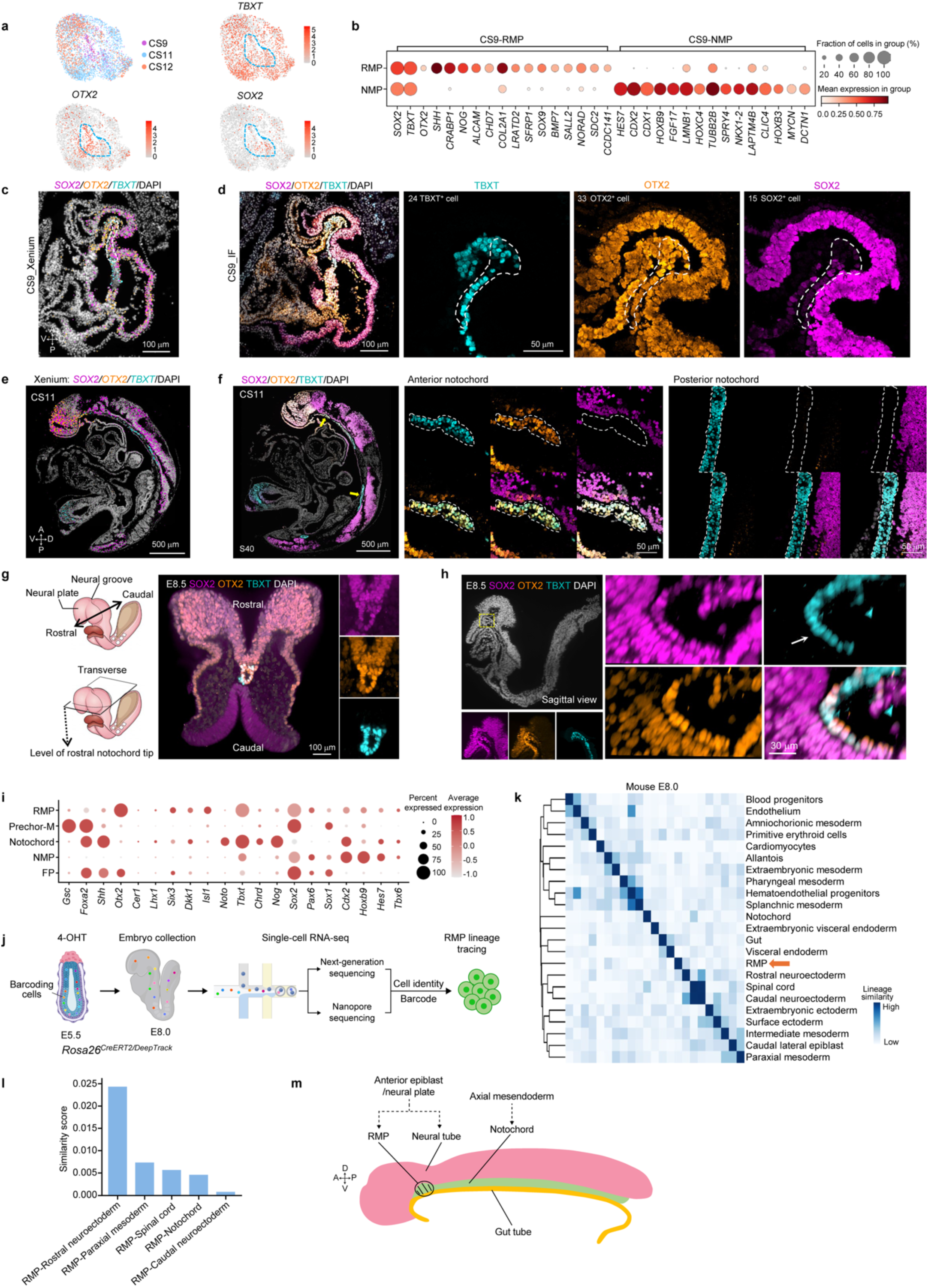
A molecularly distinct axial progenitor population emerges at the rostral tissue domain (a) UMAPs of axial progenitor and notochord-related populations from CS9 (magenta), CS11 (blue), and CS12 (orange), with expression of *TBXT*, *SOX2*, and *OTX2* shown, respectively. (b) Dot plot comparing the top differentially expressed genes between RMP cells and NMP cells at CS9. Dot size represents the percentage of cells expressing each gene; dot color indicates average expression level. (c) Representative Xenium images showing *SOX2*, *TBXT*, and *OTX2* expression at CS9. Scale bar, 100 µm. (d) Immunofluorescence staining for SOX2, TBXT, and OTX2 on a CS9 section adjacent to the Xenium section shown in (c). Scale bars as indicated. (e) Representative Xenium section with *SOX2, TBXT*, and *OTX2* expression is shown at CS11. (f) Immunofluorescence staining for SOX2, TBXT, and OTX2 on the CS11 sections adjacent to the Xenium sections, respectively. Notochord is highlighted by dotted circles. Yellow arrowheads indicate rostral and caudal notochord, respectively. Scale bars as indicated. (g) Whole-mount immunofluorescence staining of a mouse E8.5 embryo for SOX2, OTX2, and TBXT. (left) Schematic of an E8.5 mouse embryo in lateral view, indicating the neural groove, neural plate, and rostro-caudal axis. Representative transverse section with SOX2, OTX2, and TBXT staining are shown. Sectioning planes, orientations, and viewing angles are indicated. Scale bar, 100 µm. (h) Sagittal section of a mouse E8.5 embryo with the staining of SOX2, TBXT and OTX2. Yellow dashed box indicates zoom-in area in the right panel. Scale bars are 30 µm. (i) Dot plot benchmarking the molecular identity of RMPs against reference populations from published mouse E8.0-9.0 scRNA-seq data (Cao, S. et al). Dot size represents the percentage of cells expressing each gene, and dot color indicates the average expression level (log-normalized counts) in expressing cells. RMP, rostral midline progenitor; prechor-M, prechordal mesoderm; NMP, neuromesodermal progenitor; FP, floor plate. (j) Schematic of the experimental workflow. 4-hydroxytamoxifen (4-OHT) was administered at embryonic day (E) 5.5. Embryos were collected at E8.0, dissociated, and processed for single-cell RNA sequencing (scRNA-seq). Sequencing libraries were split for short-read next-generation sequencing and long-read Nanopore sequencing. (k) Hierarchically clustered heatmap of pairwise lineage-coupling scores based on shared lineage barcodes among E8.0 cell types (rows and columns are in identical order). RMP cells (orange arrow) show a close lineage relationship to rostral neuroectoderm. Color intensity indicates the strength of the relationship; the dark diagonal reflects self-comparisons. (l) Bar graph showing lineage similarity score of RMP and associated cell types caculated based on data shown in (k). (m) A sagittal view of embryo illustrating the anatomical relationship of the RMP to the neural tube, notochord, and gut tube, together with its lineage origin. A, anterior; P, posterior; D, dorsal; V, ventral.

Adjacent to rostral notochord, we identified a population of *SOX2⁺ TBXT⁺ OTX2⁺* (STO^+^) cells in the CS9 embryo (**Figs. 4a-c**). Immunofluorescence confirmed the presence of this population, located ventral to the anterior neural plate (**Fig. 4d**). Because SOX2 and TBXT co-expression is characteristic of NMPs, we first tested whether this population corresponds to NMPs. However, NMPs localize to the posterior, whereas STO⁺ cells occupy a rostral midline position. Direct comparison showed that the STO^+^ population expressed low or undetectable levels of the NMP-associated genes *CDX1, CDX2, HES7, HOXB9*, and *NKX1-2* (**Fig. 4b**). At CS11 and CS12, TBXT and OTX2 co-expression was retained while SOX2 was absent (**Figs. 4e-f, Supplementary Fig. 9b**), indicating that the STO^+^ population had acquired a unique cell state upon differentiation.

To place this population (hereafter referred to as rostral midline progenitors, RMP) in the context of midline signaling centers, we investigated mouse embryo, and identified an analogous *Sox2^+^Tbxt^+^Otx2^+^*RMP-like population in rostral tissue in E8.0 and E8.5 embryos (**Figs. 4g-h**, **Supplementary Figs. 9c-f**). Transverse sections revealed tissue continuity between the neural plate and the SOX2^+^TBXT^+^OTX2^+^ cells (**Supplementary Fig. 9e**). At the intersection where the neural plate, prechordal plate, and foregut meet, OTX2⁺SOX2⁺ neural plate cells co-expressed TBXT, whereas cells located further ventrally showed reduced SOX2 but high OTX2 and TBXT expression (**Fig. 4g, Movie 4**). The neural plate dorsal to the prechordal or notochordal plate showed a mediolateral transition in cell morphology, from pseudostratified columnar at the lateral margins to stratified cuboidal at the midline, where they directly contacted the underlying prechordal or notochordal plate (**Supplementary Fig. 9h**). The integrity of the basement membrane at the interface was assessed by checking LAMININ, SOX2, and TBXT. Discontinuous LAMININ staining with perforations on the ventral surface of the neural plate was observed adjacent to SOX2⁺TBXT⁺ cells, indicating remodeling basement membrane (**Supplementary Fig. 9g**). These SOX2⁺TBXT⁺ cells were N-CADHERIN^+^ (**Supplementary Fig. 9i**). Collectively, these observations suggest that the *Sox2^+^Tbxt^+^Otx2^+^*cells are segregated from the *Otx2⁺Sox2⁺* neural plate and acquire *Tbxt* expression upon delamination. Consistent with this, examination of human embryos from the Kyoto Collection revealed that cells in rostral regions corresponding to the diencephalon adopted a mesenchymal morphology (**Supplementary Fig. 9j**). In addition, we analyzed a recently published mouse scRNA-seq atlas and consistently identified the RMP population^69^. We compared RMP cells against prechordal mesoderm, notochord, rostral neural plate, and NMP. RMP cells expressed higher levels of *Otx2, Sox2, Six3*, and *Pax6*, suggesting a strong anterior neuroectoderm identity, but expressed undetectable or lower levels for *Gsc, Foxa2,* and *Shh*, a combination that distinguishes them from prechordal mesoderm (**Fig. 4i**). Thus, although RMP cells share markers with axial midline populations, their transcriptional profile is distinct from both prechordal mesoderm and NMPs.

To directly probe the lineage origin of RMPs, we analyzed our in vivo clonal lineage tracing data from a related study using a Cre-loxP based barcoding mouse model, termed DeepTrack^70^. This approach enables individual cell fates to be tracked via indelible cellular barcodes and allows joint profiling of gene expression in single cells. Barcodes were induced with 4-hydroxytamoxifen (4-OHT) at the pre-gastrulation stage (E5.5) in *Rosa26^CreERT2/DeepTrack^* mouse embryos and then assessed at E8.0 using scRNA-seq (**Fig. 4j**). Clonal analysis revealed that *Sox2⁺Tbxt⁺Otx2⁺* RMP cells are clonally related to rostral neuroectoderm, consistent with transcriptomic comparisons and supporting a development identity for RMPs that is distinct from axial mesoderm (**Figs. 4k-l**).

Together, our data identify a rostral midline progenitor population that is distinct from both prechordal mesoderm and NMPs in transcriptional profile, lineage relationship, and anatomical location (**Fig. 4m**). We show that this population appears to be conserved in mice and, has not previously been resolved in humans.

### In vitro derivation of RMP-like cells and their inductive capacity

We then asked whether STO^+^ RMP cells could be recapitulated from human pluripotent stem cells (hPSCs) in vitro. We noted that the OTX2^+^ anterior neural plate exhibited higher cell density, higher Hippo pathway activity, and higher Wnt signaling activity, but lower BMP signaling activity, relative to the posterior neural tube (**Supplementary Figs. 10a-b**). To approximate the high cell density and epithelial geometry of the anterior neural plate, hPSCs were micropatterned on rectangular substrates^71^ and maintained in pluripotency-supporting medium (mTeSR Plus). After 5 days, a heterogeneous cell population comprising SOX2^+^OTX2^+^ and SOX2^+^OTX2^-^ subpopulations emerged in the culture (data not show). Further culture in N2 medium at day 5-7, yielded a nearly homogeneous anterior epiblast/neural plate-like cells expressing SOX2, OTX2 and PAX6 (**Figs. 5a-b**). scRNA-seq further confirmed that the majority of cells co-expressed *SOX2*, *OTX2*, *HESX1, PAX6*, and *SIX3,* resembling anterior epiblast/neural plate, whereas some cells transcriptionally resembled neural crest and non-neural ectoderm (**Fig**. **5c**). Upon short induction with media containing WNT agonist (CHIR99021), bFGF, and BMP antagonist (LDN-193189) for 18h, these anterior epiblast/neural plate-like cells upregulated TBXT while retaining SOX2 and OTX2 expression, recapitulating RMP-like cells (**Supplementary Fig. 10c**).

**Figure 5.**
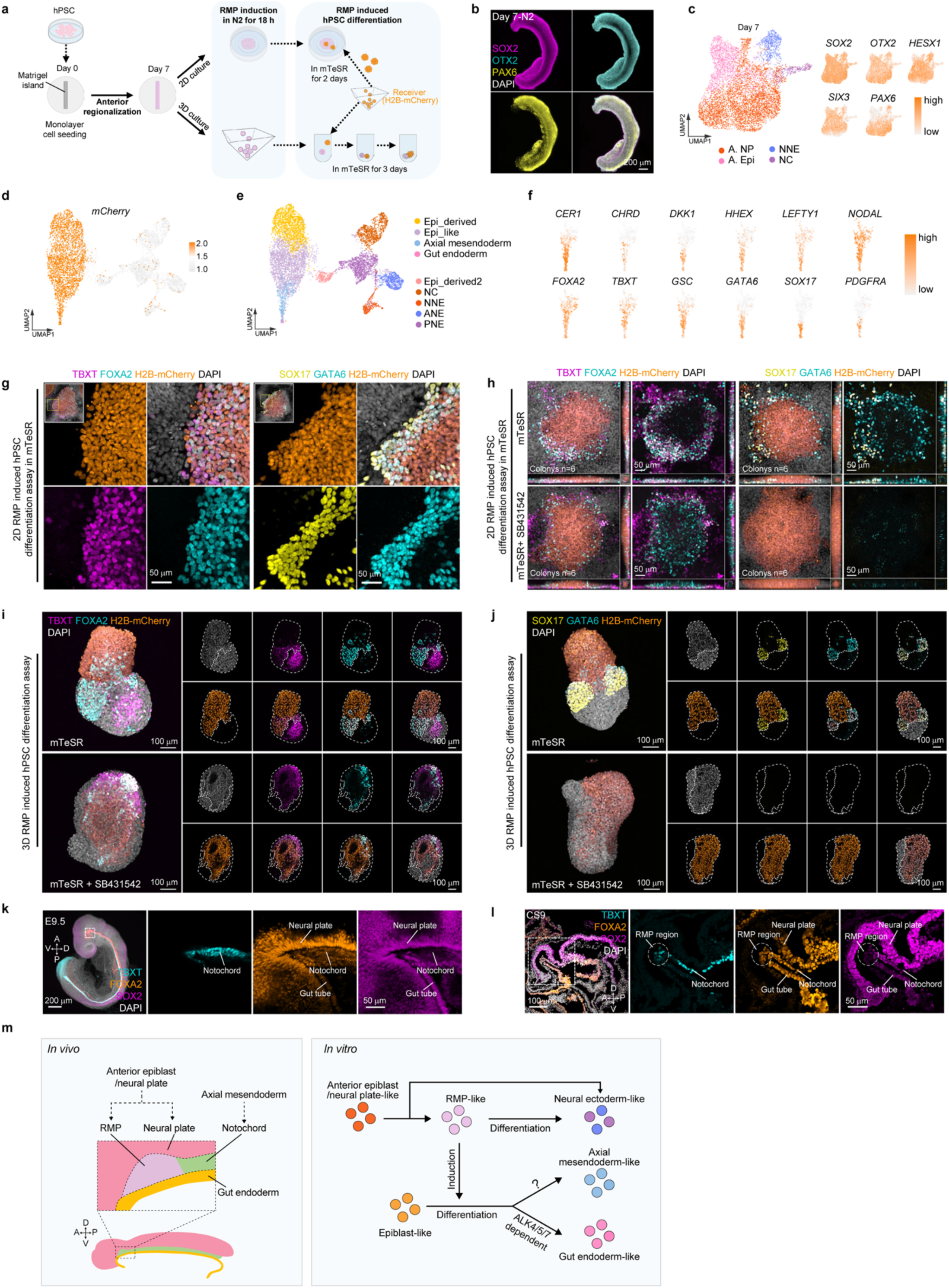
In vitro derivation of RMP-like cells and their inductive capacity (a) Schematic of the experimental workflow for in vitro differentiation of RMP-like cells from human pluripotent stem cells (hPSCs) and co-culture of RMP-like cells with hPSCs in 2D and 3D. (b) Immunofluorescence staining of anterior neural plate/epiblast-like cells at day 7 for SOX2 (magenta), OTX2 (cyan), PAX6 (yellow), and DAPI. Scale bar, 200 μm. (c) scRNA-seq of micropatterned hPSC-derived cells at day 7, before induction. Left, UMAP colored by cell type. Clusters were annotated using canonical marker genes. Most cells resemble anterior epiblast/neural plate, and smaller clusters resemble neural crest and non-neural ectoderm. Right, feature plots of anterior-associated genes OTX2, HESX1, PAX6, SIX3, SOX2 on the same UMAP. (d) UMAP of all cells from the 2-day co-culture, colored by mCherry transcript level. H2B-mCherry⁺ (with any detectable mCherry transcript) recipient hPSC-derived cells (orange) and unlabeled RMP-like feeder-derived cells (gray) occupy largely separate regions of the embedding. The small number of mCherry⁺ cells within the feeder region were teated as presumed doublets. (e) UMAP of scRNA-seq from the 2-day co-culture of RMP-like feeder cells and H2B-mCherry hPSC spheres. Clusters were annotated using canonical markers. (f) Marker gene expression in mCherry⁺ recipient cells from the co-culture (re-embedded subset of the UMAP in d), supporting the cluster annotations in (e): axial mesendoderm markers (TBXT, FOXA2, GSC, CER1, and the organizer-associated genes CHRD and DKK1), gut endoderm markers (HHEX, GATA6, SOX17), NODAL signaling components (NODAL, LEFTY1), and the mesoderm marker PDGFRA. Color scale, normalized expression (low to high). (g) Immunofluorescence staining of 2D co-cultures of H2B-mCherry-labeled hPSC spheres (right of the dashed box) on RMP-like cell feeders (left of the dashed box), after 2 days; single-channel and merged images are shown for FOXA2 and TBXT. Recipient hPSC spheres express FOXA2 and TBXT, while the feeder RMP-like cells have lost TBXT, indicating exit from the initial state. Scale bar, 50 μm. (h) Immunofluorescence staining of 2D co-cultures of RMP-like cell feeders with H2B-mCherry-labeled hPSC spheres, after 2 days, in the absence or presence of the ALK4/5/7 inhibitor SB431542. Recipient spheres in the co-culture express TBXT, FOXA2, GATA6, and SOX17; with SB431542, TBXT and FOXA2 are retained but SOX17 and GATA6 are absent, indicating that ALK4/5/7 signaling is required for the endoderm component of the induction. Orthogonal (X-Z and Y-Z) projections of an individual mCherry⁺ colony are shown. Scale bar, 50 μm. (**i-j**) Immunofluorescence staining of RMP-like cell with hPSC in 3D co-culture for TBXT, FOXA2, GATA6, and SOX17 in the presence of the NODAL inhibitor SB431542. Dashed line indicates sample edge and mCherry^+^ cell region. Scale bar, 100 μm. (k) Whole-mount and optical sections of mouse E9.5 embryos stained for SOX2, FOXA2, and TBXT. Dashed box indicates zoom-in area in right panel. Scale bars as indicated. (l) Sagittal section of a CS9 human embryo stained for SOX2, TBXT, and FOXA2. Dashed box indicates zoom-in area in right panel. Scale bars as indicated. (m) Schematic summary. In vivo, the RMP is directly apposed to the neural plate, notochord, and gut endoderm. In vitro, anterior epiblast/ neural plate-like cells convert into RMP-like cells upon CLF (<u>C</u>HIR99021, <u>L</u>DN193189, b<u>F</u>GF) treatment; RMP-like cells differentiate into neural ectoderm-like cells, while epiblast-like cells can be induced and differentiated into axial mesendoderm-like and gut endoderm-like cells by RMP-like cells. A. NP, anterior neural plate; A. Epi, anterior epiblast; NNE, non-neural ectoderm; NC, neural crest; ANE, anterior neural ectoderm; PNE, posterior neural ectoderm. A, anterior; P, posterior; D, dorsal; V, ventral.

Next, we asked whether RMP-like cells possess lineage inductive capacity. The in vitro SOX2^+^OTX2^+^TBXT^+^ cells were plated as a feeder layer, onto which hPSCs spheres carrying a knock-in mCherry-H2B reporter were seeded (**Fig. 5a**). After 2 days in co-culture, the recipient hPSC spheres robustly expressed TBXT, FOXA2, GATA6, GATA4, and SOX17, but were negative for NKX6-1 (**Figs. 5g, Supplementary Fig. 10d**), whereas the bulk of feeder cells lost SOX2, TBXT, and OTX2 expression, suggesting exit from their initial fate. scRNA-seq of the co-culture confirmed that recipient cells (mCherry^+^) comprised gut endoderm-like (*FOXA2^+^, HHEX^+^, OTX2^+^*) and axial mesendodermal-like cells (*TBXT^+^, FOXA2^+^, CER1^+^, GSC^+^*), while the RMP-like cells-derived cells adopted ectodermal fates consistent with their in vivo counterparts (**Figs. 5d-f**). The same induction was observed in 3D co-culture, in which RMP-like cells induced FOXA2⁺TBXT⁺ axial mesendoderm and GATA6⁺SOX17⁺ gut endoderm in hPSC spheroids (**Figs. 5i-j, Supplementary Fig. 10f**). Induced fates were radially arranged, peripherally in closer proximity to the RMP-like cells-derived cells (**Figs. 5g-h, Supplementary Figs. 10f-g**), mirroring the rostral body organization in vivo (**Figs. 5k-l**).

Transcriptomic analysis of in vitro RMP-like cells revealed expression of ligands and co-factors of the NODAL, BMP, WNT, FGF, and SHH pathways (**Supplementary Fig. 10h**). To identify the signal responsible, we inhibited each pathway in the co-culture. Blockade of TGF-beta/Nodal signaling with the ALK4/5/7 inhibitor SB431542 abolished the induction of gut endoderm (GATA6⁺SOX17⁺) but not of axial mesendoderm (FOXA2⁺TBXT⁺), in both 2D and 3D formats (**Figs. 5h-j, Supplementary Fig. 10e**). Conversely, inhibition of FGF/MEK (PD0325901), BMP (LDN-193189), WNT (IWR-1), or SHH (Sonidegib) signaling did not prevent the induction of FOXA2⁺TBXT⁺ cells (**Supplementary Fig. 10g**). Thus, RMP-mediated induction comprises two signaling-dependent arms: a ALK4/5/7-dependent arm driving gut endoderm, and a ALK4/5/7-independent arm driving axial mesendoderm that is not accounted for by FGF, BMP, WNT, or SHH signaling alone.

Together, these results demonstrate that RMP-like cells can be generated in vitro through sequential induction by mimicking the in vivo signaling environment. Furthermore, RMP-like cells function as a signaling source for neighboring epiblast cells, inducing gut endoderm through ALK4/5/7-mediated signaling and axial mesendoderm through a ALK4/5/7-independent route (**Fig. 5m**).

## DISCUSSION

In this study, we present a spatial transcriptomic atlas of early human organogenesis spanning post-conception weeks 3-5, including one of the first 3D transcriptomic maps of the human CS11 and CS12 stages. We identify and localize organ progenitor populations within their native anatomical context. 3D reconstruction of embryonic structures and tissue-specific gene enrichment provides a spatial framework for charting the transcriptional programs and signaling pathways underlying early organogenesis.

Embryonic midline structures pattern the overlying neural tube and somites. At the rostral tip of embryonic midline, we identify a distinct cell population, termed rostral midline progenitor (RMP). RMP cells localize adjacent to the rostral notochord/prechordal plate, a developmentally pivotal yet technically challenging region that has remained underexplored due to its small population size^72^. Direct transcriptomic comparison showed that RMP cells are transcriptionally closest to rostral neuroectoderm and distinct from both prechordal mesoderm and NMPs. Of note, the prechordal mesoderm signature is defined almost entirely from mouse and chick studies, in which the tissue was isolated and its organizing activity tested by grafting and ex vivo culture^73–81^. Human prechordal mesoderm, by contrast, has never been isolated or functionally tested. Whether RMP cells correspond to previously described anterior definitive endoderm, axial mesoderm or axial mesendoderm^82,83^, remains an open question. Our clonal barcoding analysis in mouse indicates that RMP cells are clonally related to rostral neuroectoderm, supporting a lineage relationship distinct from axial mesoderm, which contributes to head structures including the extraocular muscles and peri-ocular mesenchyme^77,81^. Functionally, RMP-like cells induce gut endoderm and axial mesendoderm in neighboring epiblast cells, both non-neural germ layer fates, in contrast to the neural-inducing activity characterized for prechordal mesoderm^73,74,76,77,79,81^. Specifically, induction of gut endoderm by RMP-like cells requires TGF-beta/Nodal signaling, whereas induction of axial mesendoderm is TGF-beta/Nodal-independent. The signals that drive the axial-mesendoderm induction remains to be identified. The identification of RMP cells in mouse provides an experimentally tractable system for dissecting their molecular features and their relationship with rostral neural plate.

Stem cell-based embryo models (SCBEMs) are widely used to study human development, with many recapitulating peri-implantation stages and some having the potential to progress to gastrulation and organogenesis^84–100^. However, even SCBEMs that mimic gastrulation^95,101–105^ and early organogenesis^71,106–110^, generally lack functional inductive signaling centers, such as the notochord and floor plate, and fail to faithfully model the rostral body. Natural embryo references are essential for both inspiring and benchmarking those models. Leveraging this spatial atlas, we further provide a rationale for the directed differentiation of hPSCs toward the anterior epiblast, neural plate, and RMP-like state in vitro. In co-culture, RMP-like cells induce gut endoderm and axial mesendoderm fates in neighboring epiblast-like cells, enabling in vitro modeling of the rostral body and inclusion of the axial mesodermal^17^. We compared several recently published organogenesis models with our atlas. Models of cardiogenesis and somitogenesis showed relatively high transcriptional similarity to their in vivo counterparts, with the cardiac model performing particularly well across CS11 and CS12 stages, though certain specialized cell types, including pacemaker cells, were not yet represented^109,111^. The neural tube model exhibited moderate overall similarity^110^. Cell types corresponding to the spinal cord and neural crest were reasonably well captured, while rostral brain fates and signaling-center populations (floor plate, roof plate) appeared less faithfully recapitulated, suggesting that rostral and midline patterning remains an area for continued refinement. These comparisons exemplify the utility of our atlas as a benchmark for guiding the iterative improvement of SCBEMs (**Supplementary Fig. 11**).

## LIMITATION

We acknowledge several limitations of this study. Due to the scarcity of human embryonic material, each Carnegie stage is represented by a single specimen, and the spatial sections do not encompass the entire embryo, with lateral tissues paticularly underrepresented. For reasons of scope and space, several hallmark structures of this developmental window, notably the optic vesicle and the otic placode, were not analyzed here, and extraembryonic cell types were excluded from analysis. The upper limb bud appears at approximately CS12, however, their lateral position places them at the edges of the sagittal section series, so they are underrepresented in the profiled sections. The in vitro RMP-like cells provide a tractable functional proxy, but remain an approximation of the in vivo population, and may not fully reproduce their in vivo state. The lineage origin of the RMP population is supported by clonal barcoding in mice and by its transcriptional proximity to rostral neuroectoderm, but direct lineage tracing in the human embryo is not feasible. Further work will be required to delineate its developmental potential and to test its inductive activity in vivo.

## ACKNOWLEDGEMENT

We thank Dr. Olivier Pourquié for providing the mCherry-H2B knock-in NCRM1 hiPSC line and Dr. Ali H. Brivanlou for providing the RUES2 hESC line. We thank Dr. Kathryn Cheah, Dr. Joshua Mark Brickman, Dr. Bernadette S. de Bakker, Dr. Patrick Tam, Dr. Tiago Rito, Dr. Aryeh Warmflash, Dr. Jun Wu, and Dr. Alfonso Martinez Arias for valuable discussions and insights. We acknowledge the support received from the Strategic Priority Research Program of the Chinese Academy of Sciences (XDB0820000, XDB1350300), the National Natural Science Foundation of China (32570954, 32541005, 82525028, 32450525, 82270123), the National Key R&D Program of China (2022YFA1105700), Zhejiang Provincial Natural Science Foundation of China (LR25C070001), and Key R&D Program of Zhejiang (2024SSYS00XX).

## AUTHOR CONTRIBUTIONS

L.L. and L. Y. conceptualized the project. **Y.Y., Q.C., H.R.,** T.L., N.C., X.L., F.H., R.L., N.Z., and X.L. collected and processed the embryos and performed data acquisition. **Q.C.,** T.L., X.L., and L.L. staged the embryos. **G.T.**, **Y.W.**, **H.R.**, and **M.L.** performed bioinformatic analyses, 3D reconstruction, and data presentation. **M.L.** performed mouse embryo collection and data acquisition. **Y.Y.** and Y.L. generated hPSC models and performed micropattern experiments. S.Y. and G.S. acquired human embryo sections and histological images. X.L. and R.L. prepared illustration of embryonic structures. C.G. generated, and J.J. and W.P. analyzed the Polylox barcoding data. **L.L., Y.Y., G.T., Y.W.,** and **M. L.** interpreted the data and drafted the manuscript. L.Y., T.L., W.P., and G.S. proofread and edited the manuscript. L.L., L.Y. T.L., W.K. supervised the project. and acquired funding.

## METHODS

### Ethics statement

Human embryos used in this study were obtained from elective terminations of pregnancy from healthy donors at the Chongqing Maternal and Child Health Hospital (approval no. 2024-048) with written informed consent. All embryo samples were discarded as clinical material. The decision to terminate the pregnancy was made prior to and independently of the decision to donate tissue for research. The study protocol was reviewed and approved by the institutional ethics committee, and sample collection followed the principle of minimal collection.

Human pluripotent stem cell experiments were performed at Westlake University and were approved by the Embryonic Stem Cell Research Oversight Committee. All experiments were conducted in accordance with the Guidelines for Stem Cell Research and Clinical Translation released by the International Society for Stem Cell Research (ISSCR, 2025 update). The culture duration of hPSCs derived cutlure was carefully limited to the minimum period necessary to examine cell fate decisions. All experiments on mice were conducted in accordance with ethical review by the Westlake University Institutional Animal Care and Use Committee. All procedures were conducted in accordance with relevant institutional and regulatory guidelines for animal care and use.

### Human embryo sample collection and preparation

Embryos used for Xenium and Visium HD analyses were morphologically intact. Developmental stages were assigned based on embryo size, somite number, anatomical features, and gestational age, and were classified as CS9 (late), CS11, and CS12 according to the Carnegie staging system. The samples were obtained and transported into cold PBS to remove surface blood. Subsequently, the embryos were placed in 4% PFA overnight at 4 ℃, followed by PBS washing, dehydration, permeabilization, paraffin exchange, and finally embedded in paraffin. Continuous sections were obtained sagittally, 5 µm in thickness, with intervals of 5 µm, 15 µm or 20 µm, were used in samples CS9, CS11, and CS12.

### Spatial transcriptomics using Visium HD

The prepared paraffin blocks (containing tissue samples fixed in 10% NBF after isolation, dehydrated, and embedded in paraffin) were sectioned on a microtome (four sections each for CS9, CS11 and CS12). Sections were floated on a water bath and mounted onto designated glass slides. Slides were then baked at 45°C for 3 h in an oven, followed by deparaffinization and rehydration: xylene was used to remove paraffin, and a descending ethanol gradient was applied to rehydrate the sections. After rehydration, hematoxylin and eosin (H&E) staining was performed to visualize morphological features, and bright-field microscopy was used for imaging and scanning. Subsequently, overnight probe hybridization was conducted. The probes consisted of two segments: a left-hand side (LHS) probe and a right-hand side (RHS) probe. The LHS was linked to Read 2, whereas the RHS carried a Poly(A) structure. Both probes hybridized to adjacent regions of the same target gene sequence. After hybridization, non-specific probes were removed by washing, and a ligase was used to join the two probe segments into a complete probe, followed by tissue washing with buffer. Slides with hybridized and ligated probes, together with tissue capture chips, were loaded into the CytAssist instrument according to the manufacturer’s instructions. RNAse treatment was applied to degrade the RNA template strand bound by the probes. Under the action of a permeabilization enzyme, the DNA probes were released in situ from cells and transferred vertically to the spots of a gene expression chip, where the Poly(A) tail on the RHS was captured by oligonucleotide primers on the chip. Probe extension was then performed via PCR, and the products were eluted from the capture chip into EP tubes. During library construction, qPCR was first used to determine the optimal amplification cycle number. Sample index PCR (SI-PCR) was subsequently performed to introduce dual-index adapters, followed by magnetic bead purification and library quality control. Quantification was carried out using Qubit 3.0. Libraries that passed quality control were sequenced on the Illumina NovaSeq 6000 platform with PE150 reads. Raw sequencing data were processed using the 10x Genomics Space Ranger HD pipeline (v4.0), which performed alignment to the human (*Homo_sapiens*) GRCh38 reference genome, filtering, and quantification, yielding gene expression matrices at bin resolutions of 2, 8, and 16 μm. These data were integrated with corresponding high-resolution fluorescence images for spatial registration, mapping the gene expression matrix onto the spatial coordinates of the tissue section.

### Spatial transcriptomics analysis using 10x Xenium

Formalin-fixed, paraffin-embedded (FFPE) tissue sections (5 μm thick) were mounted onto Xenium slides. Deparaffinization and permeabilization were then performed according to the 10x Genomics user guides (CG000578, CG000580). Probe hybridization, ligation, and amplification steps were carried out following the 10x Genomics protocol (CG000582). Briefly, overnight probe hybridization was performed at 50 °C using a custom gene panel (final probe concentration: 10 nM) targeting approximately 5,000 mRNAs. Subsequent stringent washes removed unbound probes, followed by a 2 h ligation reaction at 37 °C. Rolling circle amplification (RCA) primers were annealed during this step. Ligated, circularized probes were enzymatically amplified for 2 h at 37 °C to generate multiple copies of gene-specific barcoded signals per RNA binding event. After washing, background fluorescence was chemically quenched. Finally, the slide was assembled into an imaging cassette and loaded into the Xenium Analyzer (v3.1.0.0) instrument for image acquisition as per the manufacturer’s instructions (CG000584). Raw image data were processed by the integrated Xenium Analyzer pipeline for decoding and expression quantification, yielding a cell-by-feature expression matrix. The resulting subcellular-resolution spatial transcriptomics data were interactively explored and visualized using the 10x Genomics Xenium Explorer software (v3.1.0.4), including inspection of cell segmentation, spatial expression visualization, and clustering and downstream analysis based on the expression matrix.

### Cell and nucleus segmentation

Cell and nucleus segmentation were performed using the integrated Xenium Onboard Analysis pipeline. Nuclei were identified from DAPI-stained morphology images using the 10x Genomics custom neural-network-based nucleus segmentation algorithm, which incorporates information from the three-dimensional DAPI image stack. Cell boundaries were subsequently inferred by isotropic expansion of the segmented nuclear boundaries by up to 5 μm or until an adjacent cell boundary was encountered. Overlapping regions generated during nuclear expansion were resolved using an algorithm conceptually similar to Voronoi tessellation. Decoded transcripts located within the resulting cell boundaries were then assigned to individual cells to generate the cell-by-feature expression matrix.

### Spatial data processing and clustering

To maximize the preservation of spatial tissue architecture and ensure data quality, we implemented distinct quality control procedures tailored to the different sequencing platforms. For Visium HD data, the Load10X_Spatial function in Seurat (v5.2.1) was used to load the data and create Seurat objects. Subsequent filtering removed cells expressing fewer than 50 genes (min. features = 50) or with mitochondrial gene expression exceeding 5%, while retaining genes detected in at least 3 cells (min. cells = 3). For Xenium data, analysis objects were constructed using the LoadXenium function in Seurat (v5.2.1), retaining only cells with a nucleus area greater than zero. To integrate all datasets, slice data from different samples were first merged and underwent uniform preprocessing: each sample was normalized independently, and 2,000 highly variable genes (HVGs) were identified within each sample. Gene expression data were then scaled and subjected to principal component analysis (PCA). Building on this, batch-effect correction for the integrated expression data was performed using the Harmony (v1.2.3)^112^ method. Based on the first 30 dimensions derived from Harmony-corrected data, a nearest-neighbor graph was constructed using the FindNeighbors function, followed by unsupervised clustering with the FindClusters function (resolution parameter set to 0.5). Finally, Uniform Manifold Approximation and Projection (UMAP) was performed using the same 30 dimensions for visualization of the clustering results.

### Cluster annotation

Unsupervised clustering of the integrated spatial transcriptomics data was performed using Seurat (v5.0). Based on the spatial coordinates and clustering results, differential expression analysis among clusters was conducted with the FindAllMarkers() function (using default parameters) to identify signature genes for each cluster. Differentially expressed genes (DEGs) were filtered using the following criteria: adjusted P-value (p_val_adj) <= 0.05 and log₂-transformed fold change (log₂FC) >= 0.25. The filtered genes were then sorted in descending order by log₂FC, and the top 50 genes ranked by log₂FC in each cluster were selected as candidate marker genes. Preliminary cell type annotation for each cluster was performed by cross-referencing this list of candidate marker genes with well-established, classically reported cell type marker genes. To further validate and refine spatial cell-type annotations, reference-based deconvolution was performed using matched scRNA-seq datasets. Single-cell RNA-seq data were used as reference profiles, and RCTD analysis was performed on Visium HD and Xenium datasets to estimate cell-type composition and assign spatial cell-type probabilities. The inferred cell-type compositions were compared with canonical marker expression and spatial localization patterns to support cell-type annotation.

### Comparison of *in vivo* embryonic data with human-derived *in vitro* models

We extracted the corresponding cell types from the Visium HD spatial data and subsequently integrated them with the somite organoid dataset^109^ using the FindIntegrationAnchors and IntegrateData functions in Seurat. Following the same workflow, we separately projected the human neural tube model dataset^110^ and the heart dataset^111,113^ onto the UMAP embedding derived from the Visium HD data. Using the integrated data, Spearman correlation coefficients were calculated for 2,000 highly variable genes via the cor function in R (v 4.4.3).

### Spatial region-specific differential gene expression analysis

To identify differentially expressed genes across distinct spatial regions, we first selected and extracted the barcodes corresponding to target areas based on spatial coordinates. Using these barcodes, a scanpy (v1.11.4) analysis object was constructed. Differential gene expression analysis was then performed with the sc.tl.rank_genes_groups function, and the results were visualized using sc.pl.rank_genes_groups_dotplot.

### scRNA-seq library preparation and sequencing

For single-cell RNA sequencing, the collected tissue was washed twice with PBS buffer and then subjected to enzymatic digestion. The digestion protocol involved initial treatment with 1 mg/ml collagenase type IV and 1 mg/ml DNase I, followed by digestion with Trypsin-EDTA and additional DNase I at 37°C for 5 minutes. Subsequently, the single-cell suspension was filtered through a 70 μm cell strainer, washed with D-PBS, and resuspended in D-PBS containing 0.4% BSA at approximately 1000 cells/μL for sequencing. During library preparation, cells were diluted to about 5000 cells per well and loaded onto a Chromium Next GEM Chip G along with the master mix according to the 10X Genomics Chromium Next GEM Single Cell 3’ Reagent Kit v3.1 protocol. Following GEM-RT purification, the subsequent steps were carried out as per the manufacturer’s instructions.

### scRNA data processing

Raw sequencing data were processed using Cell Ranger (v5.0.1) and aligned to the GRCh38 reference genome and annotation with default parameters. Subsequent analyses, including preprocessing, normalization, identification of variable features, data scaling, dimensionality reduction, clustering, and identification of key marker genes, were primarily conducted using Seurat. To remove apoptotic or dead cells, we retained cells with a detected gene count (nFeature_RNA) greater than 200 and a mitochondrial gene percentage (percent.mito) below 20%. Key marker genes for each cluster were primarily identified using the FindAllMarkers function.

### 3D reconstruction

Serial sections were reconstructed into three-dimensional embryo models. Adjacent tissue sections were registered using the SuperPoint-SuperGlue framework based on image feature matching^36,37^. SuperPoint was used to detect keypoints and extract local feature descriptors, with a maximum of 1,024 keypoints per image, a keypoint confidence threshold of 0.005, and a non-maximum suppression radius of 4 pixels. Feature matching between adjacent sections was subsequently performed using the pretrained SuperGlue network with the indoor weights, 20 Sinkhorn iterations, and a matching confidence threshold of 0.2. The resulting matched feature points were used for spatial alignment of the corresponding sections. Gene expression matrices obtained from Xenium spatial transcriptomics sequencing together with their corresponding x-y spatial coordinates, multiple two-dimensional slices were integrated. Slice registration was performed using the Partial alignment method implemented in Spateo (version 0.0.0)^114^, with the parameter partial_robust_level set to 50 to enhance robustness against local variations. Following slice registration, three-dimensional reconstruction was performed based on the aligned data, generating raw point cloud, cell mesh, and voxel-based models. For mesh reconstruction, the marching cube algorithm was applied (cs_method = “marching_cube”), and all reconstructed three-dimensional structures were subsequently visualized.

### Analysis of TF regulation

We inferred transcription factor (TF) regulatory activities from 10x Genomics Visium HD (8 μm) spatial transcriptomics data using the command-line workflow of pySCENIC (v0.12.1), following previously described methods^115^. To compare cell-type-specific regulon activities, the previously defined regulon specificity score (RSS) was applied. Based on this framework, transcription factor modules across embryonic cells spanning all developmental stages were systematically identified using an established approach. This analysis consisted of four main steps: 1) Regulon activity inference: TF regulatory activities in individual cells were inferred using the AUCell method implemented in pySCENIC; 2) Correlation analysis: Pearson correlation coefficients were calculated between the regulatory activity scores of all transcription factors; 3) Connection specificity index (CSI) calculation: For a given pair of transcription factors a and b, the CSI was defined as the proportion of transcription factors connected to a or b whose Pearson correlation coefficients with a or b were lower than the correlation coefficient between a and b; 4) Regulon module identification: Transcription factor modules were identified by hierarchical clustering of the CSI matrix using the Ward linkage method. Finally, the average regulatory activity of each transcription factor module was calculated and visualized across different cell types.

### Cell-cell communication analysis

We analyzed intercellular communication using CellPhoneDB v5.0.1^116^. Following the recommended workflow, the raw count matrix and cell-type annotations were exported from the Seurat object using the provided scripts. These files were then used as input for the statistical_analysis module with default parameters, and executed via the Linux command-line interface. Significant cell-cell interactions between key cell types were visualized using the “dot_plot” function.

### Pathway activity scoring

Pathway activity scoring and spatial trend analysis gene sets representing major developmental signaling pathways were curated, and the complete list of genes included in each pathway is provided in Supplementary Table 5. Pathway activity at the single-cell level was evaluated using the irGSEA framework. Gene-set transcriptional scores were independently calculated using AUCell, UCell, singscore, single-sample gene set enrichment analysis (ssGSEA), JASMINE, and Viper, and concordance across scoring methods was assessed using robust rank aggregation (RRA). OTX2⁺ and OTX2⁻ cells were defined as cells with OTX2 expression above zero and equal to zero, respectively. The pathway-associated transcriptional scores were computed per cell with AUCell using predefined pathway gene sets within each cell’s expression ranking. Scores were compared with a two-sided Wilcoxon rank-sum test, which compares the overall rank distributions of the two groups and not only their medians

### Rostrocaudal and dorsoventral spatial trend analysis

For rostral-to-caudal (R-C) analysis, a continuous curved spatial coordinate was constructed from the aligned two-dimensional single-cell spatial coordinates. A k-nearest-neighbor (kNN) graph was generated based on spatial proximity, with each cell connected to its 12 nearest neighbors and Euclidean distances used as edge weights. Rostral and caudal spatial anchors were defined at the two ends of the embryo, and shortest-path distances from each cell to both anchors were calculated using *Dijkstra*’s algorithm. For cell i, the R–C coordinate was defined as:

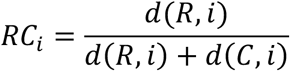

Where *d*(*R*, *i*) and *d*(*C*, *i*) represent the shortest-path distances to the rostral and caudal anchors, respectively. The resulting coordinate, termed RC_curve, was scaled from 0 (rostral) to 1 (caudal). Cells were ordered according to RC_curve and divided into 60 equal-width bins. Spatial trends in pathway activity were fitted using generalized additive models (GAMs) with shrinkage cubic regression splines, with the basis dimension set to k=8. In parallel, cells from all slices within the same developmental stage were combined and fitted using the same model to derive an overall R–C trend representing the shared spatial pattern across slices.

For dorsal–ventral (D–V) analysis, representative cell types exhibiting spatial continuity along the D–V axis were selected. For each selected cell type, representative gene sets corresponding to developmental signaling pathways were analyzed using the same irGSEA framework to characterize pathway activity patterns across the D–V spatial organization.

### Cross-platform assessment of spatial resolution using RCTD

To compare cell-type compositional resolution across single-cell RNA sequencing, Xenium, Visium HD, and a published spatial transcriptomic dataset, all datasets were evaluated using a unified single-cell reference and the Robust Cell Type Decomposition (RCTD) framework. A multistage single-cell RNA-seq dataset was used to construct the common reference, while the entire CS12-7 sample was held out as an independent test dataset and was not included in reference construction. Analyses were restricted to genes shared across the single-cell, Xenium, Visium HD, and published spatial datasets to ensure that all platforms were evaluated within the same feature space.

For the held-out CS12-7 dataset, each experimentally profiled single cell was treated as an independent query. Xenium and Visium HD datasets were analyzed separately by spatial slice, whereas the published spatial dataset was analyzed by tissue section. All queries were deconvolved against the same single-cell reference using RCTD in both doublet and full modes. The major RCTD parameters were set to gene_cutoff = 1.25 × 10⁻⁴, fc_cutoff = 0.5, gene_cutoff_reg = 2 × 10⁻⁴, fc_cutoff_reg = 0.75, and UMI_min = 100. For multi-cell-type decomposition, the maximum number of cell types allowed per spatial location was set to four (MAX_MULTI_TYPES = 4). For the doublet mode, the proportions of locations classified as singlet, doublet_certain, doublet_uncertain, or reject were calculated to characterize the overall degree of cell-type compositional mixing. For the full mode, RCTD-derived cell-type weights were normalized for each location, and four quantitative metrics were calculated: Top1 weight, Top1–Top2 margin, normalized Shannon entropy, and the effective number of inferred cell types. Top1 weight was defined as the largest normalized cell-type weight at each location, whereas the Top1–Top2 margin was defined as the difference between the largest and second-largest weights.

For normalized cell-type weights *p_k_*, Shannon entropy was calculated as

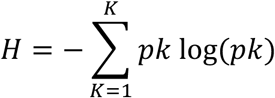

and normalized by the total number of reference cell types K:

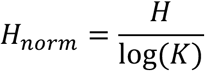

Higher normalized entropy indicates a more dispersed cell-type weight distribution and greater compositional ambiguity. The effective number of inferred cell types was calculated as

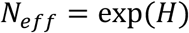

which represents the equivalent number of equally weighted cell types corresponding to the observed weight distribution. Because CS12-7 consisted of experimentally profiled single cells that were excluded from reference construction, its RCTD weight distributions were used as an empirical single-cell benchmark. Xenium, Visium HD, and the published spatial dataset were subsequently compared with this benchmark based on Top1 weight, Top1–Top2 margin, normalized entropy, effective number of inferred cell types, and doublet-mode classification composition to assess their relative cell-type compositional resolution.

### 3D embryo region selection

Spatial transcriptomics data generated using the 10x Genomics Xenium platform were registered to a three-dimensional (3D) embryonic model and stored in AnnData format^114^. Following registration, a global 3D point cloud comprising all spatially resolved cells was constructed, together with surface mesh and voxel-based embryo representations to support anatomical visualization and spatial context interpretation.

Using the Spateo framework, we performed interactive 3D rectangular cropping within the point cloud space to manually define regions of interest (ROIs)^114,117^. ROI delineation was guided by established embryonic anatomical landmarks and further validated by the regional composition of stage-specific cell types. For Carnegie stages CS9, CS11, and CS12, each embryo was subdivided along the anterior– posterior axis into anterior (A), middle (M), and posterior (P) regions. Anterior regions were characterized by enrichment of neural plate– and cranial mesoderm–derived populations, including neural tube subtypes, neural crest, and cardiac progenitors. Middle regions exhibited hallmark trunk-associated lineages such as paraxial mesoderm derivatives, neuromesodermal progenitors, and elongating neural tube populations, whereas posterior regions were enriched for tailbud, hindgut, allantois, and extraembryonic interface cell types. Each spatially defined ROI was exported as an independent VTK model. Subsequently, 3D coordinates and cell barcodes were extracted from each ROI and intersected with those in the fully registered Xenium AnnData object, retaining only shared cells. Spatial coordinates were appended to the subset expression matrices, and region-specific AnnData objects were generated for all downstream spatial and transcriptomic analyses.

### Stamp Fabrication

In this study, stamps were fabricated using a photocuring 3D printing method. Stamps with micropatterned structures were produced through stamp model design, model slicing, photocuring 3D printing, stamp cleaning and curing, and stamp sterilization. The detailed procedures are described as follows.

#### Stamp model design

AutoCAD software was used for stamp design and three-dimensional modeling. The stamp consisted of a base, a handle structure, and embossed patterns designed on the base surface. The specific micropatterns could be customized according to experimental requirements. In this study, the micropatterns were rectangular, with a length of 3 mm and a width of 0.3 mm.

#### Model slicing

The model designed in AutoCAD was exported in STL format and imported into slicing software (CHITUBOX) for slicing and support generation to prepare for subsequent 3D printing. Transparent resin was used for 3D printing via photocuring.

#### Photocuring 3D printing of stamps

The sliced PRZ file was transferred to the 3D printer using a USB drive. High-precision transparent photosensitive resin was added to the resin tank, and the printing platform was installed before starting the printing process. After printing was completed, the printing platform was removed, and the stamp was carefully detached from the platform using a scraper. (Note: As the resin is slightly toxic, the printer should be placed in a fume hood, and masks and gloves should be worn during operation.)

#### Stamp cleaning and curing

The printed stamps were cleaned using Gaopuda model cleaning solution and ultrasonicated at 40 kHz for 3 min. After discarding the cleaning solution, the stamps were ultrasonicated in absolute ethanol for 1 min, then in 75% ethanol for 1 min. During the process, the stamps were fully immersed in the cleaning solution or ethanol to ensure thorough cleaning. The printing platform and scraper were cleaned with 75% ethanol. (Note: As the resin is slightly toxic and volatile in alcohol, masks and gloves should be worn throughout the procedure.) After cleaning, the supports on the stamps were removed before curing. The stamps were then exposed to ultraviolet light for 1 min on each side to ensure complete curing.

#### Stamp sterilization

The cured stamps were disinfected with 75% ethanol and further exposed to ultraviolet light for more than 30 min in a biosafety cabinet to complete sterilization.

### Human stem cell lines and culture

RUES2 (female hESC line), RUES2-GLR (female hESC line) and *HES7*-Achilles-CAG-H2B-mCherry reporter male hiPSC line (NCMR1 background) were used in this study. All cell lines were cultured on feeder-free Matrigel-coated culture plates in mTeSR^TM^ Plus medium. To perform routine passaging, Dispase was used to digest the colonies without dissociating them into single cells. Cells were incubated in Dispase for 5 min before removal and were followed by the addition of 1 mL fresh mTeSR^TM^ Plus medium. The colonies were then lifted from the culture plate with cell lifters, and pipetted less than 10 times to break large colonies into small clumps. 5–10% of cells were transferred to a new Matrigel-coated well for maintenance. Accumax was used for making single-cell suspensions. 10 μM ROCK inhibitor Y-27632 was supplemented in the media on the day of single cell dissociation.

### Anteriorizing of human PSCs and RMP-like cells differentiation

Rectangle resin stamps (3 mm × 0.3 mm) obtained by 3D printing were sterilized as described above. Using the aseptic stamp, dip it into 2% Matrigel, then print onto the flat bottom of a 96-well plate, and transfer to a 37℃ incubator for 15 min. Approximately 2×10^5 human ESCs were seeded onto stamp-shaped Matrigel-coated wells and cultured at 37℃ in a 5% CO2, humidified incubator for 30 min to ensure cell attachment. After that, pipetting was performed continuously to remove unattached cells, followed by fresh mTeSR^TM^ Plus media with 10 μM Y-27632, which was recorded as day 0 (D0). A rectangle-shaped cell pattern with a clear boundary was formed. Y-27632 was removed after 2 days (D2), and the fresh mTeSR^TM^ Plus media was changed daily. As the cell culture period progressed, rectangle-shaped monolayer cell patterns developed into shape-limited columnar multilayer patterns. At D5, changed to N2 media: DMEM/F12 medium supplemented with 1× N2 supplement, 1× MEM Non-Essential Amino Acids Solution, 15 mM HEPES, and 5 µM 2-mercaptoethanol for another 2 days. At this point (D7), columnar cell patterns were detached for cell identity analysis, and cells were identified as anterior progenitors. Here, 5 μM CHIR, 0.5 μM LDN193189, and 10 ng/ml bFGF were administered to induce cell differentiation. Most cells in the columnar cell pattern differentiated to SOX2^+^/OTX2^+^/TBXT^+^ cells (RMP-like cell) after 18 h.

### Co-culture of RMP-like cells with human PSC-spheres in 2D and 3D

At D7, the anterior progenitors were detached and digested into single cells by Accumax. For 2D co-culture, single-cell suspensions were seeded on a glass-bottomed plate at a density of 6×10^5/cm^2^ in N2 media with 10 μM Y-27632. For 3D co-culture, single-cell suspensions were seeded on AggreWell plates at a density of 500 cells per microwell. 24 h later, fresh N2 media with 5 μM CHIR, 0.5 μM LDN193189, and 10 ng/ml bFGF were administered to induce cell differentiation. To prepare the EPI-sphere, the H2B-mCherry reporter hiPSCs were dissociated into single-cell suspensions and seeded 500-1000 cells per microwell into AggreWell plates once RMP induction initiated. After 18 h, the anterior progenitors differentiated to STO^+^ RMP-like cells, and hiPSCs formed EPI-spheres. At this point, the differentiation medium was removed, and EPI-spheres were added to the RMP-like cell layer co-cultured for an additional 2 days in mTeSR^TM^ Plus medium. Or EPI-spheres were put together with the RMP-like cell sphere in 96-well U-bottom plate and co-cultured for an additional 3 days in mTeSR^TM^ Plus medium. For the pathway perturbation assay, small molecules (1 μM sonidegib, 1 μM SAG, 1 μM IWR-1, 0.5 μM LDN193189, 1 μM PD0325901, 10 μM SB431542, respectively) were supplied as the EPI-sphere added.

### RNAscope of FFPE slices

RNAscope was performed on FFPE CS9, CS11, and CS12 slices following the manufacturer’s protocol (RNAscope™ Multiplex Fluorescent Reagent Kit v2). Briefly, slices were baked in a dry oven for 1 h at 60℃, then deparaffinized in xylene and 100% ethanol. Incubated for 10 min with hydrogen peroxide at RT and performed target retrieval later with 1× Target Retrieval Reagent. After that, apply protease plus for 15 min at 40℃. Washing slice with fresh ddH2O and proceeded to probe hybridization at 40℃ for 2 h. After slice washing, hybridized AMP1, AMP2, and AMP3 step by step, followed by HRP signal developing and fluorophore labeling according to the probe channel (RNAscope™ Probe-Hs-TBX5-C1, RNAscope™ Probe-Hs-SOX2-C2, RNAscope™ Probe - Hs-SIX3-C3).

### Immunofluorescence staining

Samples, including cells and tissue sections, were washed with PBS before fixation with 4% paraformaldehyde (PFA) at 4°C overnight. Fixed samples were washed three times with PBS before permeabilization in PBS containing 0.5% Triton X-100 at room temperature for 1 h. Samples were then blocked with 5% normal donkey serum (NDS) in PBST (PBS + 0.5% Triton X-100) at room temperature for 1 h. Primary antibody dilution was prepared in 5% NDS in PBST. Samples were incubated in the primary antibody dilution at room temperature for 2 h or at 4°C overnight, followed by 3 PBST washes, 10 min each. After washing, samples were then transferred to a secondary antibody dilution made with 5% Donkey serum in PBST and incubated at room temperature for 1 h. After that, samples were washed 3 times for 30 min with PBST at room temperature. For whole-mount embryonic staining, the primary and secondary antibody incubation conditions were adjusted to 37°C for 24 hours. In contrast to section staining, the antibody concentration was doubled.

Following immunofluorescence staining, tissue clearing was performed by immersing the sample in CUBIC-R at room temperature for 2 hours. Imaging was conducted using the NIKON AXR confocal microscope and the Zeiss Lightsheet7 microscope.

### Image analysis

3D TIFF files were generated after reconstructing serial embryonic sections. Using Amira 3D (v2024.1, Thermo Scientific), a 3D point cloud model containing cell type annotation and spatial information was reconstructed. Individual cell type clusters were segmented using ImageJ, and subsequent 3D surface rendering was performed in Imaris (v10.1.0, Oxford) to generate three-dimensional representations of each cluster.

Microscopy files in .czi and .nd2 formats were converted to .ims format using Imaris File Converter. Further adjustments, including maximum intensity projection (MIP), pseudo-coloring, and sectioning, were carried out in Imaris.

### Clonal lineage tracing of RMPs in mice

The E8.0 single-cell lineage-tracing dataset was obtained from a separate study using the DeepTrack system described previously^70^. Briefly, *Rosa26^CreERT2/DeepTrack^* embryos were labelled at E5.5 by administration of 4-hydroxytamoxifen (4-OHT; Sigma, H6278; 0.4mg/mouse or 0.07 mg/g body weight) and collected at E8.0 for single-cell RNA sequencing. DeepTrack lineage barcodes were enriched from the corresponding cDNA libraries and resolved by Nanopore sequencing as previously described^70^. To obtain clonal barcodes, lineage barcodes were filtered based on their probability of generation (Pgen), which estimates the likelihood of a given barcode being generated independently by the recombination process^70,118^. RMP cells were defined as cells co-expressing *Sox2*, *Otx2*, and *Tbxt*. To infer lineage relationships between RMP cells and other embryonic cell types, lineage coupling was calculated using cospar.tl.fate_coupling implemented in CoSpar^119^, using only clonal barcodes retained after Pgen filtering. The resulting lineage-coupling patterns were visualized using the R package pheatmap.

## MATERIALS

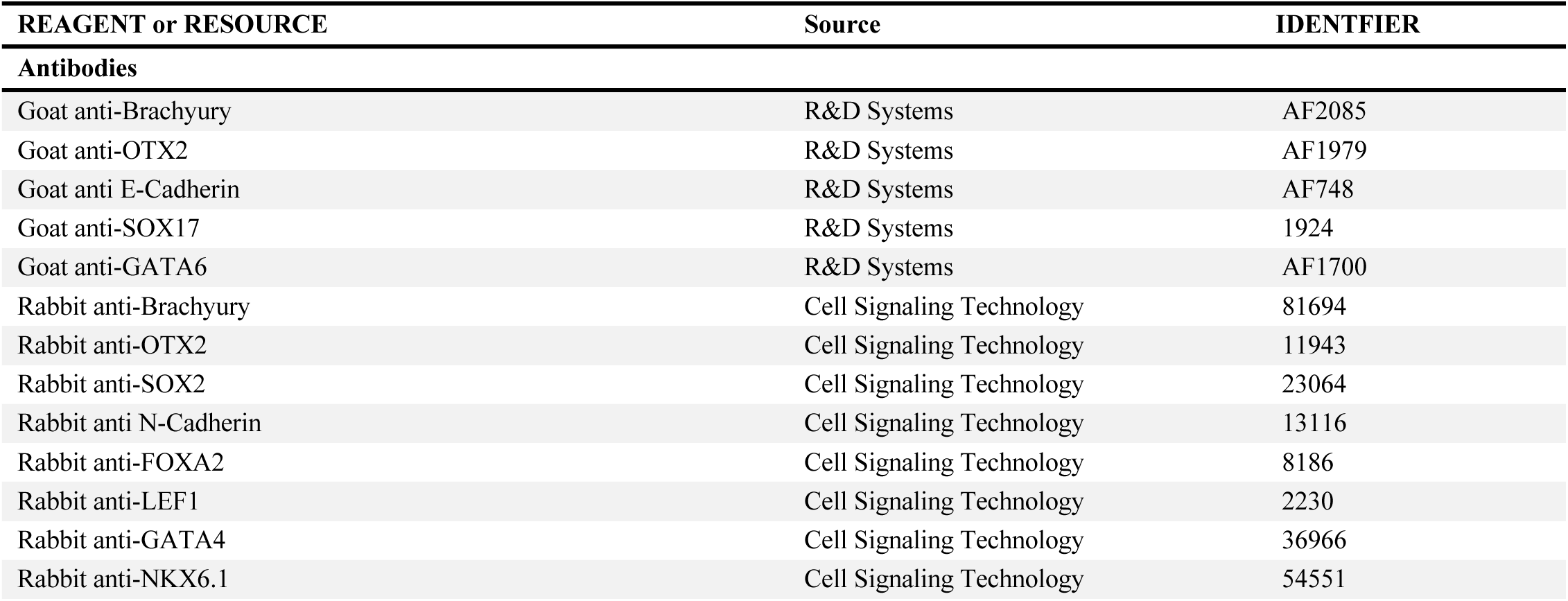
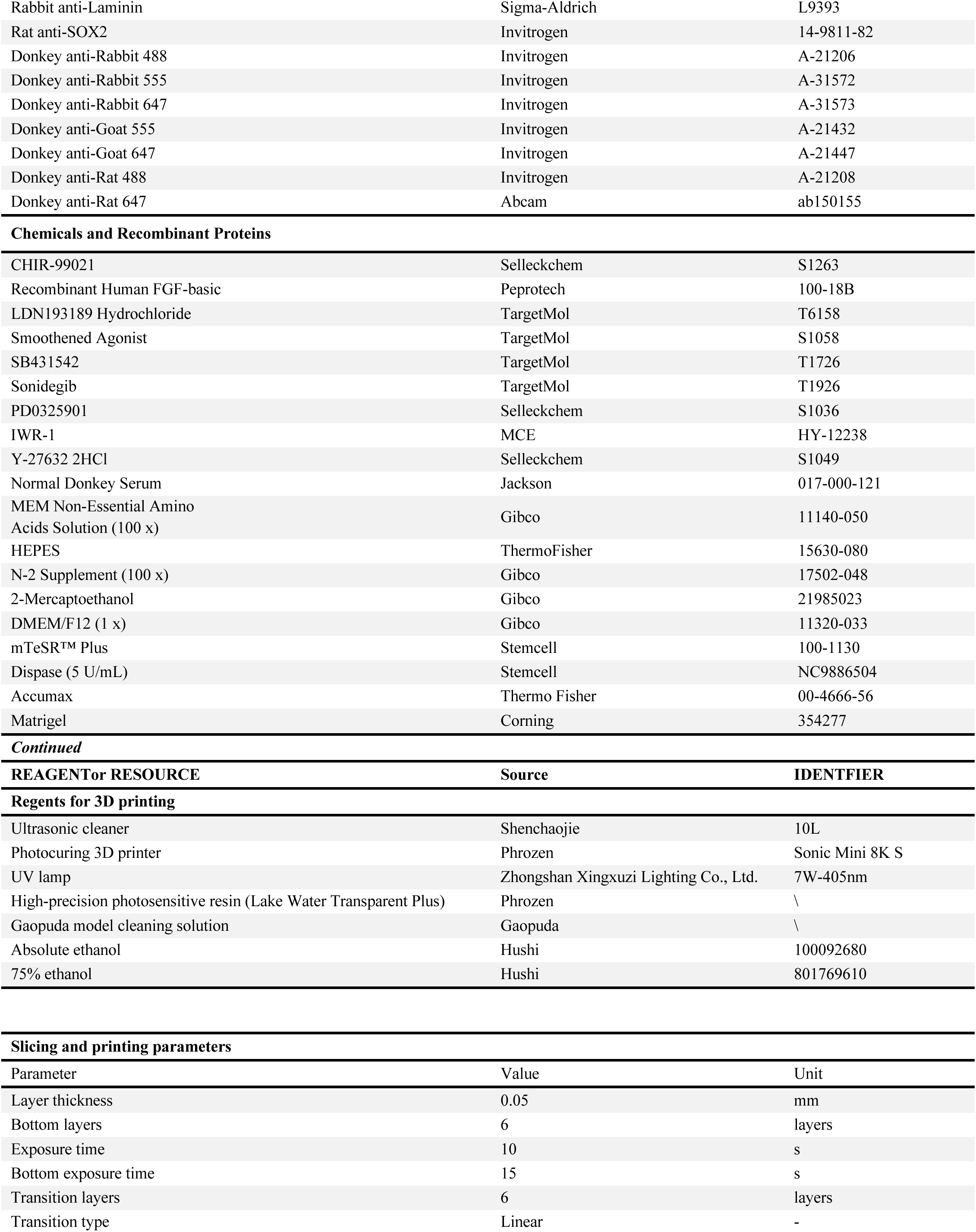
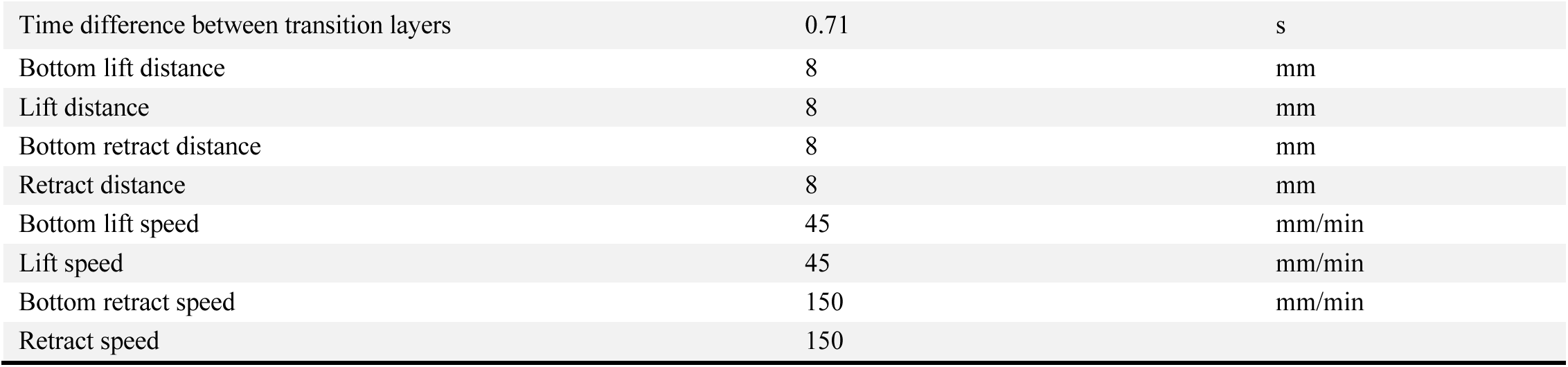

| REAGENT or RESOURCE | Source | IDENTFIER |
|---|---|---|
| **Antibodies** |  |  |
| Goat anti-Brachyury | R&D Systems | AF2085 |
| Goat anti-OTX2 | R&D Systems | AF1979 |
| Goat anti E-Cadherin | R&D Systems | AF748 |
| Goat anti-SOX17 | R&D Systems | 1924 |
| Goat anti-GATA6 | R&D Systems | AF1700 |
| Rabbit anti-Brachyury | Cell Signaling Technology | 81694 |
| Rabbit anti-OTX2 | Cell Signaling Technology | 11943 |
| Rabbit anti-SOX2 | Cell Signaling Technology | 23064 |
| Rabbit anti N-Cadherin | Cell Signaling Technology | 13116 |
| Rabbit anti-FOXA2 | Cell Signaling Technology | 8186 |
| Rabbit anti-LEF1 | Cell Signaling Technology | 2230 |
| Rabbit anti-GATA4 | Cell Signaling Technology | 36966 |
| Rabbit anti-NKX6.1 | Cell Signaling Technology | 54551 |
| Rabbit anti-Laminin | Sigma-Aldrich | L9393 |
| Rat anti-SOX2 | Invitrogen | 14-9811-82 |
| Donkey anti-Rabbit 488 | Invitrogen | A-21206 |
| Donkey anti-Rabbit 555 | Invitrogen | A-31572 |
| Donkey anti-Rabbit 647 | Invitrogen | A-31573 |
| Donkey anti-Goat 555 | Invitrogen | A-21432 |
| Donkey anti-Goat 647 | Invitrogen | A-21447 |
| Donkey anti-Rat 488 | Invitrogen | A-21208 |
| Donkey anti-Rat 647 | Abcam | ab150155 |
| **Chemicals and Recombinant Proteins** |  |  |
| CHIR-99021 | Selleckchem | S1263 |
| Recombinant Human FGF-basic | Peprotech | 100-18B |
| LDN193189 Hydrochloride | TargetMol | T6158 |
| Smoothened Agonist | TargetMol | S1058 |
| SB431542 | TargetMol | T1726 |
| Sonidegib | TargetMol | T1926 |
| PD0325901 | Selleckchem | S1036 |
| IWR-1 | MCE | HY-12238 |
| Y-27632 2HCl | Selleckchem | S1049 |
| Normal Donkey Serum | Jackson | 017-000-121 |
| MEM Non-Essential Amino Acids Solution (100 x) | Gibco | 11140-050 |
| HEPES | ThermoFisher | 15630-080 |
| N-2 Supplement (100 x) | Gibco | 17502-048 |
| 2-Mercaptoethanol | Gibco | 21985023 |
| DMEM/F12 (1 x) | Gibco | 11320-033 |
| mTeSR™ Plus | Stemcell | 100-1130 |
| Dispase (5 U/mL) | Stemcell | NC9886504 |
| Accumax | Thermo Fisher | 00-4666-56 |
| Matrigel | Corning | 354277 |
| ***Continued*** |  |  |
| **REAGENTor RESOURCE** | **Source** | **IDENTFIER** |
| **Regents for 3D printing** |  |  |
| Ultrasonic cleaner | Shenchaojie | 10L |
| Photocuring 3D printer | Phrozen | Sonic Mini 8K S |
| UV lamp | Zhongshan Xingxuzi Lighting Co., Ltd. | 7W-405nm |
| High-precision photosensitive resin (Lake Water Transparent Plus) | Phrozen | \ |
| Gaopuda model cleaning solution | Gaopuda | \ |
| Absolute ethanol | Hushi | 100092680 |
| 75% ethanol | Hushi | 801769610 |
| **Slicing and printing parameters** |  |  |
| Parameter | Value | Unit |
| Layer thickness | 0.05 | mm |
| Bottom layers | 6 | layers |
| Exposure time | 10 | s |
| Bottom exposure time | 15 | s |
| Transition layers | 6 | layers |
| Transition type | Linear | - |
| Time difference between transition layers | 0.71 | s |
| Bottom lift distance | 8 | mm |
| Lift distance | 8 | mm |
| Bottom retract distance | 8 | mm |
| Retract distance | 8 | mm |
| Bottom lift speed | 45 | mm/min |
| Lift speed | 45 | mm/min |
| Bottom retract speed | 150 | mm/min |
| Retract speed | 150 |  |

---

**Supplementary Figure 1.**
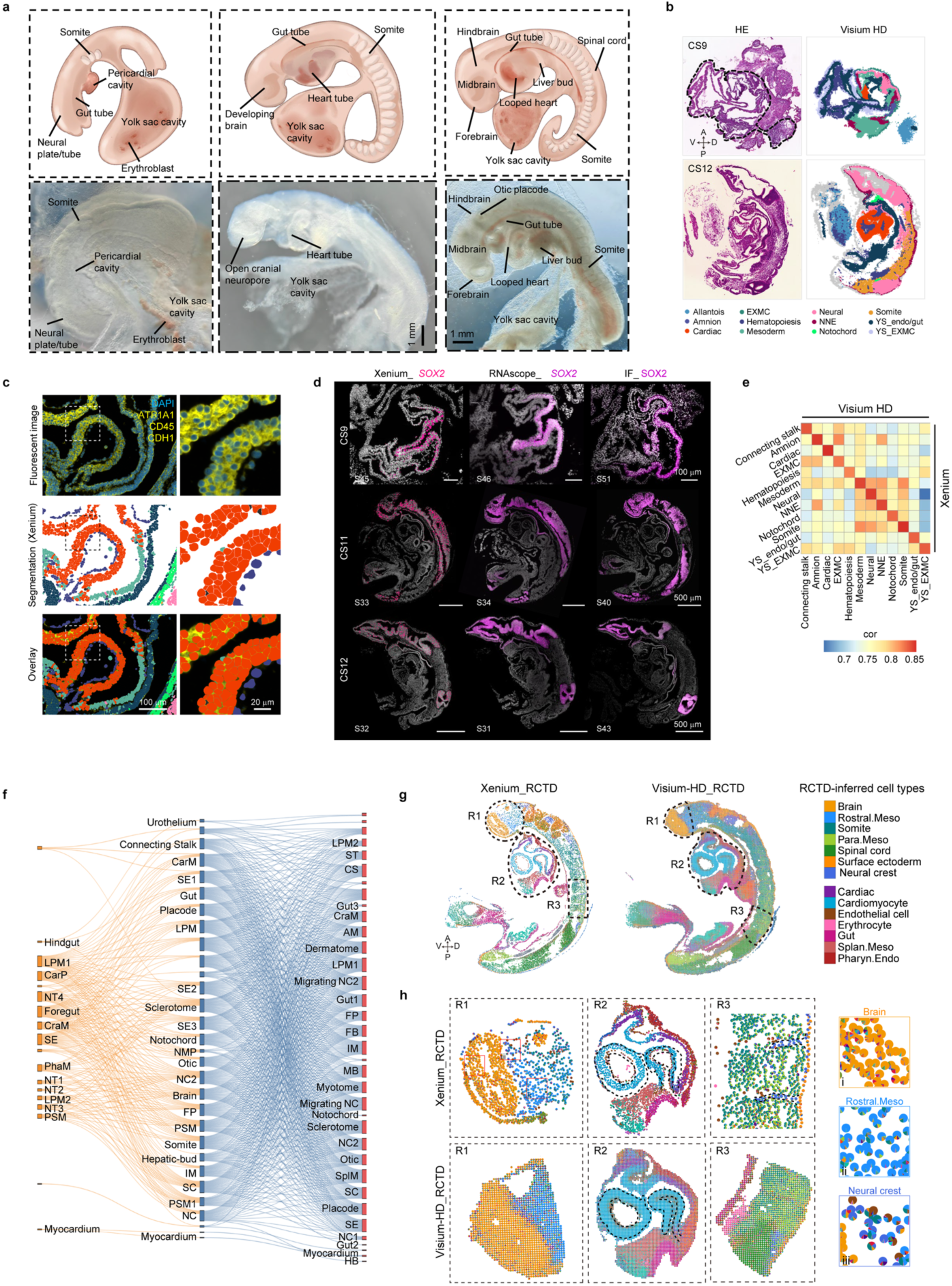
Spatial transcriptomic profiling of human CS9, CS11, and CS12 embryos (associated with Figure 1). (a) Schematics and representative images of embryos analyzed in this study, with key anatomical structures labeled. Scale bars as indicated. (b) Representative Visium HD and Xenium sections from CS9 and CS12 with corresponding HE staining. The embryo and yolk sac are indicated by a dotted circle at CS9. Placental tissue is not shown the Visium HD visualization. (c) Cell segmentation based on Xenium fluorescence imaging. DAPI marks nuclei; ATP1A1, CD45, and CDH1 (E-CADHERIN) mark cell membranes. Scale bar as indicated. (d) Representative sections of CS11 examined by Xenium, RNAscope, and immunofluorescence. Slice number (s) as indicated. Scale bars as indicated. (e) Concordance between Visium HD and Xenium platforms assessed by pseudo-bulk Spearman correlation of gene expression profiles. (f) Correspondence of cell types identified across CS9, CS11, and CS12, shown as a lineage relationship diagram. (g) Xenium and Visium HD spatial maps after deconvolution with robust cell-type decomposition (RCTD). Full-section cell-type map with three regions (R1-R3, dashed outlines) selected for magnification. Cell types are color-coded. (h) Magnification area for Xenium maps at single-cell resolution, each segmented cell is rendered as a pie chart depicting its cell-type assignment in the magnified views. NNE, non-neural ectoderm; EXMC, extraembryonic mesoderm; YS_endo/gut, yolk sac endoderm and primitive gut; YS_EXMC, yolk sac extraembryonic mesoderm; NMP, neuromesodermal progenitor; PSM, presomitic mesoderm; LPM, lateral plate mesoderm; NT, neural tube; SE, surface ectoderm; NC, neural crest; FP, floor plate; FB, forebrain; MB, midbrain; HB, hindbrain; Otic, otic placode; SC, spinal cord; CraM, craniofacial mesoderm; CarP, cardiac progenitor; CarM, cardiac mesoderm; PhaM, pharyngeal mesoderm; ST, septum transversum; AM, anterior mesoderm; IM, intermediate mesoderm; SplM, splanchnic mesoderm. A, anterior; P, posterior; D, dorsal; V, ventral.

**Supplementary Figure 2.**
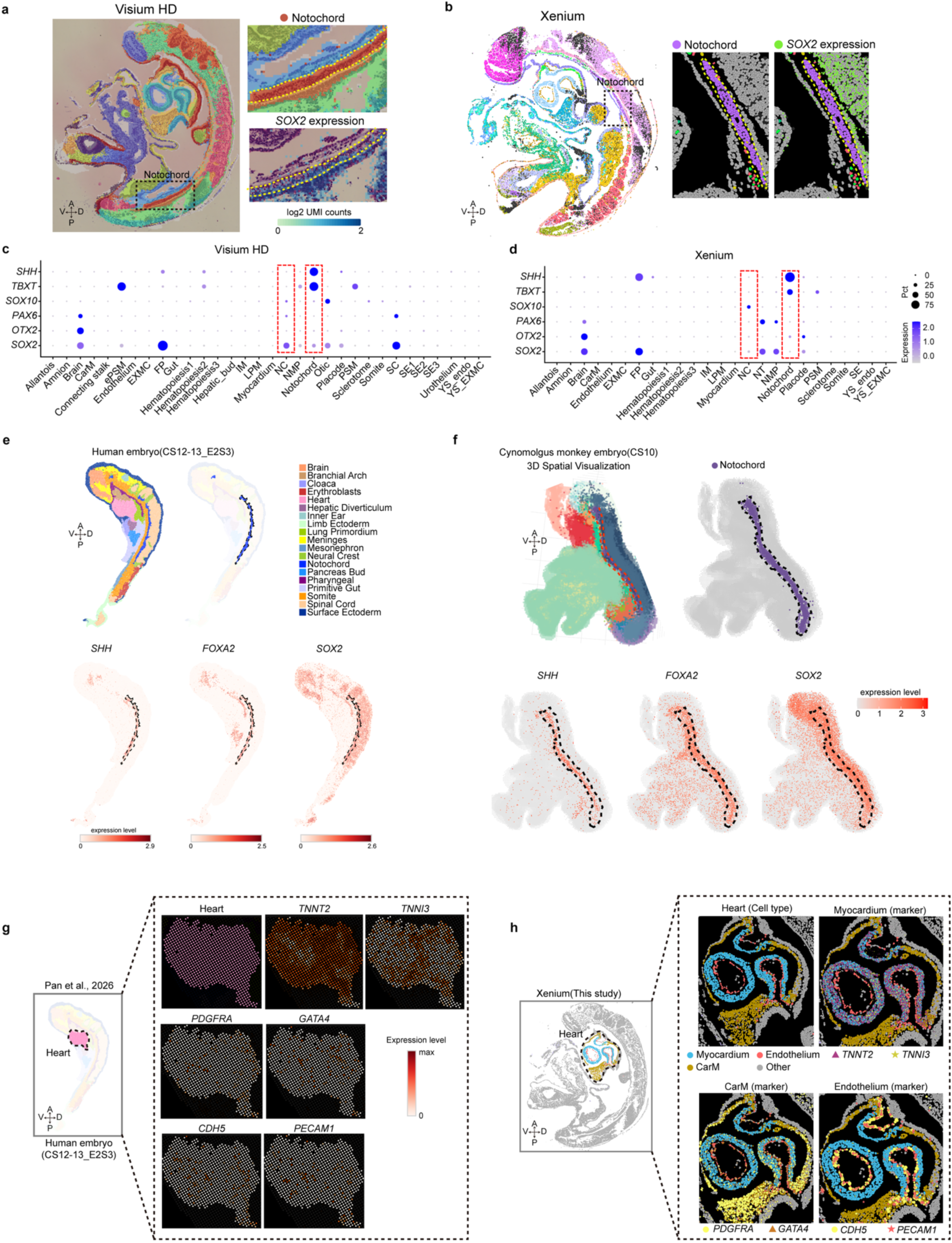
Signal blending and cell type misclassification in capture-based sequencing platforms Visium-HD and Stereo-seq (associated with Figure 1). (**a-b**) Spatial clustering and cell-type annotation of the Xenium and Visium HD dataset, respectively. The left panel shows the overall spatial distribution of the embryo, with dashed boxes indicating representative regions. Enlarged views of notochord and the spatial expression of *SOX2* in the notochord and adjacent tissues. (**c-d**) Dot plots of marker gene expression for the clusters compared as in (a, b); red dashed boxes highlight *SOX2* expression in notochord and neural crest. (e) Spatial distribution of the notochord and representative gene expression in a published human embryo spatial transcriptomics dataset (Pan et al. 2026). The upper panel shows the original tissue annotation and the location of the notochord, whereas the lower panels show the spatial expression of *SHH*, *FOXA2*, and *SOX2*. Dashed lines indicate the notochord region. (f) Spatial distribution of the notochord and representative gene expression in a three-dimensional spatial transcriptomics dataset of a cynomolgus monkey embryo (Tan et al. 2026). The upper panels show the overall spatial annotation and the location of the notochord, whereas the lower panels show the spatial expression of *SHH*, *FOXA2*, and *SOX2*. Dashed lines indicate the notochord region for comparison with the human embryo spatial dataset. Colors denote different cell types or tissue annotations, and gene-expression levels are indicated by the color scales shown in each panel. (g) Spatial expression patterns of cardiac-associated marker genes in a published human embryo spatial transcriptomic dataset (Pan et al. 2026). The left panel shows the region originally annotated as Heart, and the enlarged panels on the right show the spatial expression of the myocardial markers *TNNT2* and *TNNI3*, the mesenchymal/cardiac-associated markers *PDGFRA* and *GATA4*, and the endothelial markers *CDH5* and *PECAM1*. These markers exhibited dispersed or broadly distributed expression patterns within the Heart-annotated region, with limited spatial separation of distinct cellular populations. (h) Spatial resolution of cell types and their marker genes within the embryonic heart in the Xenium dataset generated in this study. The left panel shows the location of the heart region, and the right panels show cell-type annotations together with the spatial distributions of marker genes associated with myocardium, CarM, and endothelium. Myocardial cells are marked by *TNNT2* and *TNNI3*, CarM cells by *PDGFRA* and *GATA4*, and endothelial cells by *CDH5* and *PECAM1*. The spatial distributions of these marker genes closely correspond to their respective cell-type annotations, demonstrating the ability of Xenium to resolve distinct cellular populations and their spatial boundaries within the embryonic heart. A, anterior; P, posterior; D, dorsal; V, ventral; CarM, cardiac mesoderm.

**Supplementary Figure 3.**
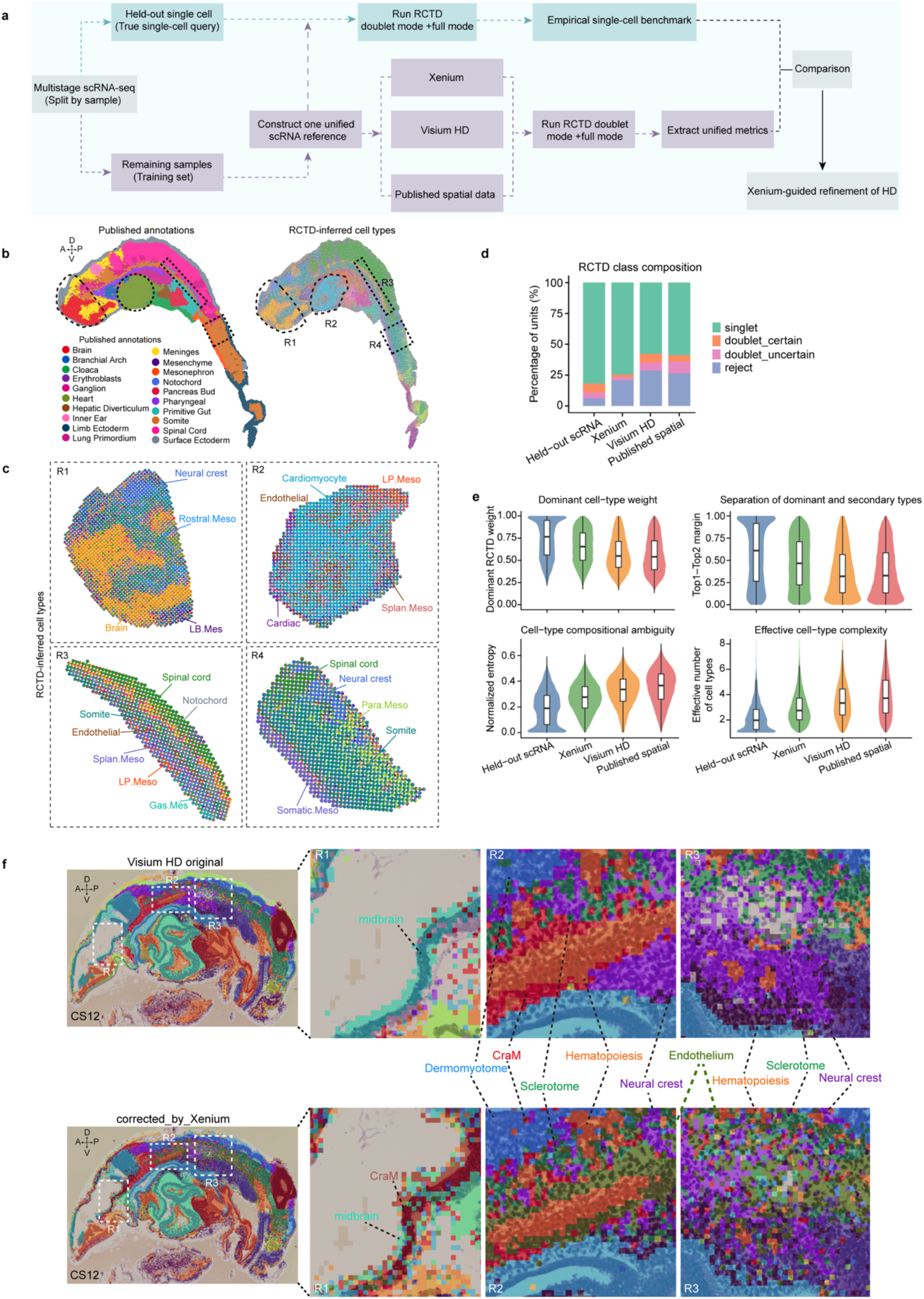
Deconvolution of Visium-HD data using Xenium data (associated with Figure 1). (a) Schematic overview of the cross-platform RCTD analysis and benchmarking framework. Published scRNA-seq datasets from matched developmental stages were first split at the sample level. One independent sample was held out as a single-cell query dataset, whereas the remaining samples were used to construct a unified scRNA-seq reference. The same reference was then applied to the held-out single-cell dataset, Xenium, Visium HD, and the published spatial transcriptomics dataset using RCTD in both doublet and full modes. Unified metrics describing cell-type composition were extracted from the RCTD results for each platform, with the independently held-out single cells serving as an empirical single-cell benchmark for cross-platform comparison and subsequent Xenium-derived reference-guided refinement of Visium HD. (**b-c**) Comparison between the original annotations and RCTD-inferred cell-type compositions in the published spatial transcriptomics dataset (Pan et al., 2016). The left panel shows the spatial region/tissue annotations reported in the original study, whereas the right panel shows cell-type compositions inferred by RCTD using the unified scRNA-seq reference. Dashed outlines indicate representative regions R1-R4, with enlarged views shown on the right to illustrate the spatial distribution and compositional heterogeneity of RCTD-inferred cell types across different anatomical regions. (d) Comparison of RCTD classification outcomes across platforms in doublet mode. The stacked bar plot shows the proportions of singlet, doublet_certain, doublet_uncertain, and reject classifications in the independently held-out single-cell dataset, Xenium, Visium HD, and the published spatial transcriptomics dataset. Singlet indicates that a spatial unit can be predominantly explained by a single cell type; doublet_certain indicates confident identification of two major cell types; doublet_uncertain indicates mixed cell-type composition with lower confidence in the second inferred cell type; and reject indicates that no reliable cell-type classification could be assigned. (e) Quantitative comparison of cell-type compositional features across platforms in RCTD full mode. Violin plots show the dominant cell-type weight, the difference between the first- and second-ranked cell-type weights (Top1-Top2 margin), normalized entropy, and effective cell-type complexity. Higher dominant cell-type weight and a larger Top1-Top2 margin indicate a more clearly defined dominant cell type, whereas higher normalized entropy and effective cell-type complexity indicate greater compositional complexity and cell-type mixing within individual spatial units. Independently held-out scRNA-seq cells were used as an empirical single-cell benchmark to assess how closely each spatial transcriptomics platform approximates single-cell compositional characteristics. (f) Xenium-guided refinement of Visium HD at CS12. The upper row shows the original spatial clustering and cell-type annotations of Visium HD, whereas the lower row shows the refined Visium HD annotations guided by high-resolution cell-type information from Xenium. The left panels show the overall spatial distribution of the CS12 embryo, with dashed boxes indicating representative regions R1, R2, and R3; enlarged views of these regions are shown on the right. Colors indicate different cell types. A, anterior; P, posterior; D, dorsal; V, ventral; CraM, craniofacial mesoderm; Rostral.Meso, rostral mesoderm; Cardiac, cardiac mesoderm; Splan.Meso, splanchnic mesoderm; LP.Meso, lateral-plate mesoderm; Para.Meso, paraxial mesoderm; Somatic.Meso, somatic mesoderm.

**Supplementary Figure 4.**
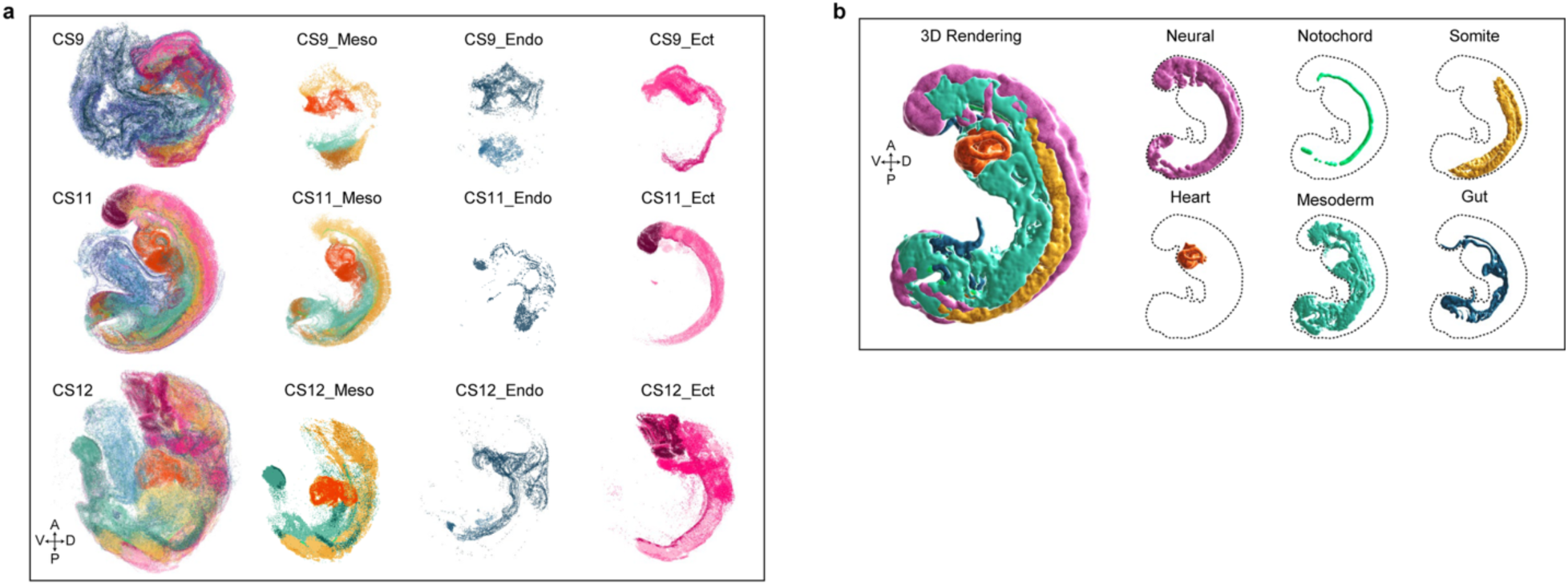
3D reconstruction of embryonic structures (associated with Figure 1) (a) Representative images of the 3D reconstruction of embryonic structures derived from Xenium in situ sequencing data. (b) Representative 3D renderings integrating spatial transcriptomics data with morphological imaging. A, anterior; P, posterior; D, dorsal; V, ventral.

**Supplementary Figure 5.**
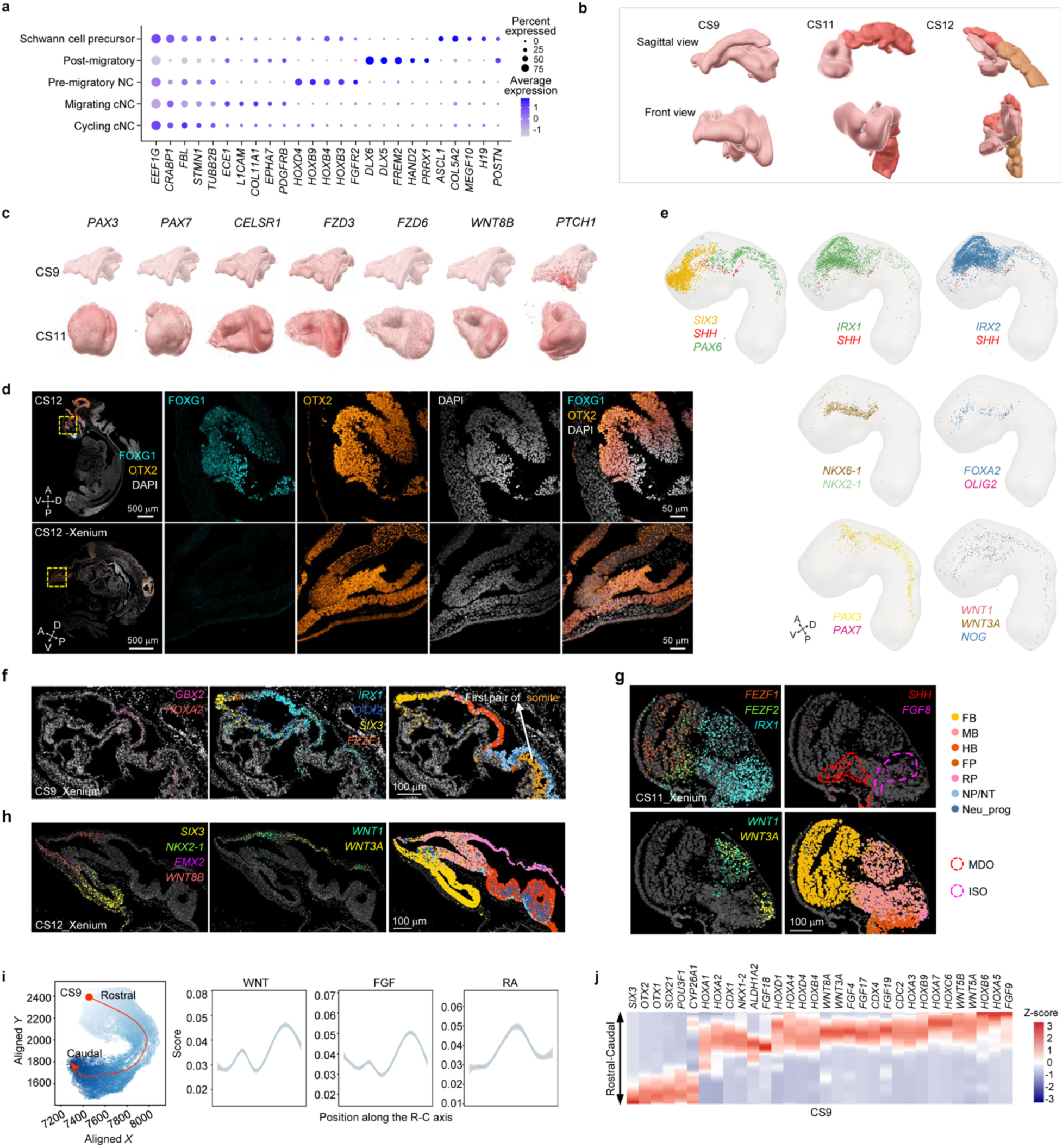
Neural crest, neural tube regionalization, and primary brain vesicle formation (associated with Figure 2). (a) Dot plot showing the top 5 genes for subtypes of neural crest. Dot size represents the percentage of cells expressing each gene, and dot color indicates the average expression level (log-normalized counts) in expressing cells. (b) 3D renderings of the neural plate and neural tube at CS9, CS11, and CS12. Renderings were reconstructed from serial sagittal sections (Methods) and are shown in sagittal (top) and front (bottom) views. At CS9, the central nervous system is an open neural plate, with elevated neural folds flanking the neural groove and a broadened rostral region corresponding to the prospective brain. At CS11, closure is under way: the rostral brain region has expanded markedly, the anterior neuropore remains visible in front view, and the caudal neural tube is still open. At CS12, the neural tube is closed along the rostrocaudal axis and visibly regionalized, with the forebrain, midbrain, and hindbrain distinguishable rostrally and the spinal cord caudally. Renderings are shown for 3D visualization only; absolute sizes are not comparable between stages. (c) 3D renderings of the neural plate and neural tube at CS9 and CS11, illustrating progressive regionalization and primary brain vesicle formation. Representative genes with regionalized expression patterns are indicated. (d) Immunofluorescence staining for FOXG1 (cyan) and OTX2 (orange) with DAPI on a CS12 section, with the corresponding Xenium spatial maps of the same markers below. The yellow-dashed box indicates the zoom-in area shown in the right panel. Scale bar, 500 µm; magnified views, 50 µm. (e) Expression of representative genes along the neural tube at CS9 displayed on 3D mesh models. (f) Representative genes displaying regionalized expression in the neural plate, neural tube, and somites at CS9. The first somite pair is indicated by an arrowhead. Scale bar, 100 µm. (g) Representative genes displaying regionalized expression in the forebrain and midbrain at CS11. The mid-diencephalic organizer (MDO) and isthmic organizer (ISO) are indicated by red and magenta dashed circles, respectively. Cell types are color-coded. Scale bar, 100 µm. (h) Representative genes displaying regionalized expression in the forebrain, midbrain, and hindbrain at CS12. Cell types are color-coded as in (**f**). Scale bar, 100 µm. (i) Signaling pathway activities along the R-C axis at CS9, quantified using irGSEA-integrated gene set pathway-associated transcriptional scores. Pathways are indicated. (j) Heatmap of differentially expressed genes along the R-C axis at CS9, ordered from rostral to caudal.

**Supplementary Figure 6.**
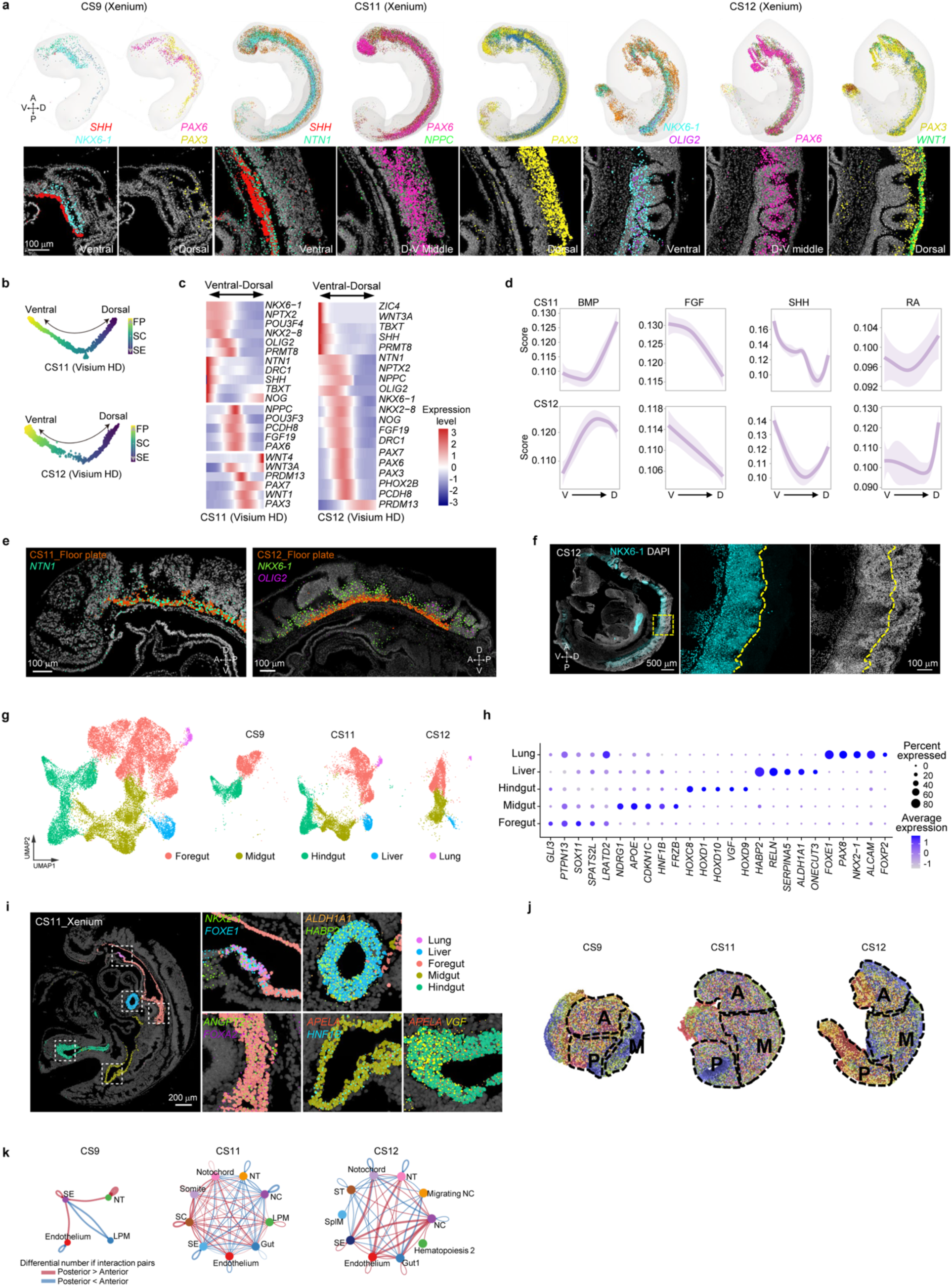
Patterning of the neural tube and endoderm (associated with Figure 2). (a) Expression of representative genes along the D-V axis at CS9, CS11, and CS12, displayed on 3D mesh models and corresponding Xenium sections. Scale bar, 100 µm. (b) Monocle2-based trajectory analysis of the D-V patterning using representative regionalized cell types. (c) Heatmap of differentially expressed genes along the D-V axis at CS11 and CS12, ordered from ventral to dorsal. (d) Signaling pathway-associated transcriptional score along the D-V axis at CS11 and CS12, quantified by integrated gene set expression scores using irGSEA. (e) Representative marker gene expression in the floor plate at CS11 (top) and CS12 (bottom). Scale bar, 100 µm. (f) Immunofluorescence staining for NKX6-1 (cyan) and DAPI at CS12. Scale bar, 500 µm. The yellow-dashed box indicates the zoom-in area shown in the right panel. The yellow-dashed line indicates the edge of NKX6-1^+^ region. Scale bar,100 µm. (g) UMAP of endoderm cells in CS9, CS11, and CS12, totally colored by five clusters, including foregut, midgut, hindgut, liver and lung. (h) Dot plot showing the top 5 genes for each identified cell cluster in (**g**). Dot size represents the percentage of cells expressing each gene, and dot color indicates the average expression level (log-normalized counts) in expressing cells. (i) Spatial map of endoderm subtypes and marker genes on the CS11 Xenium section. White-dashed boxes indicate zoom-in area shown in the right panel of individual subtypes. <u>Scale bar, 200</u> µm. (**j-k**) Spatial delineation of regions of interest selected for tissue interaction analysis across CS9, CS11, and CS12. A, anterior; P, posterior; M, middle. Regionalized tissue interactions corresponding to the regions selected in (j).

**Supplementary Figure 7.**
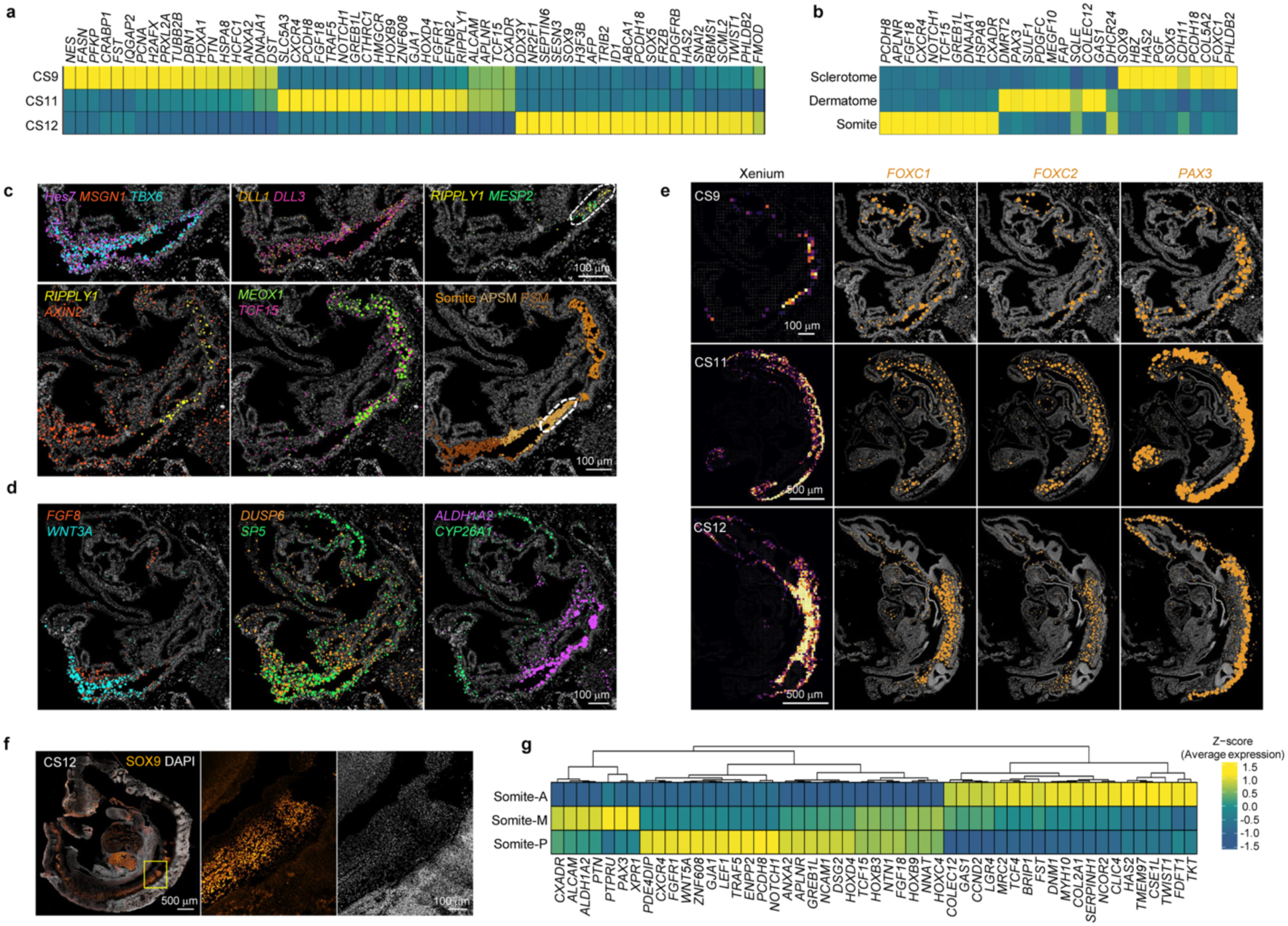
Patterning and morphogenesis of somites (associated with Figure 3) (a) Heat maps of differentially expressed genes along somite maturation at CS9, CS11, and CS12. Color-coded, yellow, high; teal, low. (b) Heat maps of differentially expressed genes among somite, dermatome, and sclerotome at CS9, CS11, and CS12. Color-coded, yellow, high; teal, low. (**c-d**) Representative Xenium sections showing expression of somitogenesis-related genes at CS9. The determination front of the newly forming somite is indicated by a dashed circle. Scale bar, 100 µm. (e) Representative Xenium sections showing the expression of *FOXC1*, *FOXC2*, and *PAX3* at CS9. (left) Composite image showing the expression density map of all three genes. (right) Individual gene expression channels as indicated. Scale bars as indicated. (f) Representative immunofluorescence staining image of the CS12-section for SOX9. Yellow box indicates the zoom-in area in right panel. Scale bars as indicated. (g) Heatmap of differentially expressed genes along thoracolumbar epithelial somites (Somite-M) and caudally positioned somites near PSM (Somite-P), and occipital/cervical somites (Somite-A).

**Supplementary Figure 8.**
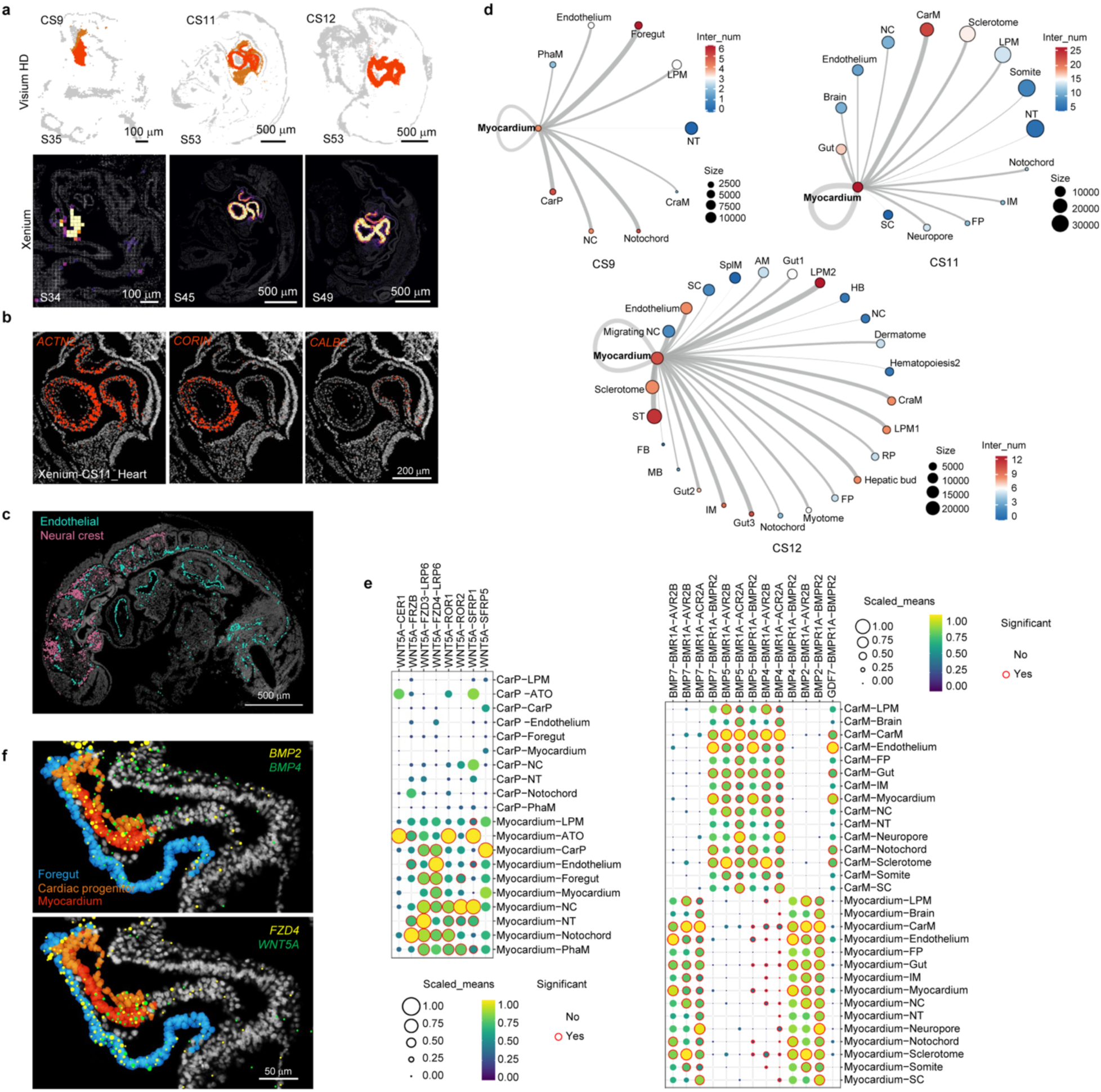
Patterning and morphogenesis of the heart (associated with Figure 3) (a) Representative Visium HD sections showing cardiac cell type (top panel) and expression gradients of cardiac genes (bottom panel), including *ACTN2*, *CALB2*, *CSRP3*, *CORIN*, *EEF1A2*, *LDB3*, *NKX2-5*, *NPPA*, *PLN*, and *TNNT2*. Scale bars as indicated. (b) Representative Xenium sections showing cardiac gene expression, corresponding to main Figure 3n-o. Scale bar is 200 µm. (c) Representative Xenium section showing endothelial and neural crest cell distributions at CS11. Scale bar, 500 µm. (d) Cell-cell communication networks involved in cardiac development at CS9, CS11, and CS12, analyzed by CellPhoneDB. (e) Dot plot showing ligand-receptor pair expression underlying indicated cell-cell communications in cardiac development at CS9 and CS11, shown in (**d**). Dot size represents the percentage of cells expressing each gene; dot color indicates average expression level (log-normalized counts). (f) Representative Xenium sections showing BMP and WNT pathway gene expression at CS9, with cell types color-coded. Scale bar, 50 µm. LPM, lateral plate mesoderm; PhaM, pharyngeal mesoderm; NT, neural tube; NC, neural crest; ATO, anterior organizer; CarP, cardiac progenitor; CarM, cardiac mesoderm; IM, intermediate mesoderm; FP, floor plate; SC, spinal cord; SplM, splanchnic mesoderm; AM, anterior mesoderm; CraM, craniofacial mesoderm; RP, roof plate; ST, septum transversum; FB, forebrain; MB, midbrain; HB, hindbrain.

**Supplementary Figure 9.**
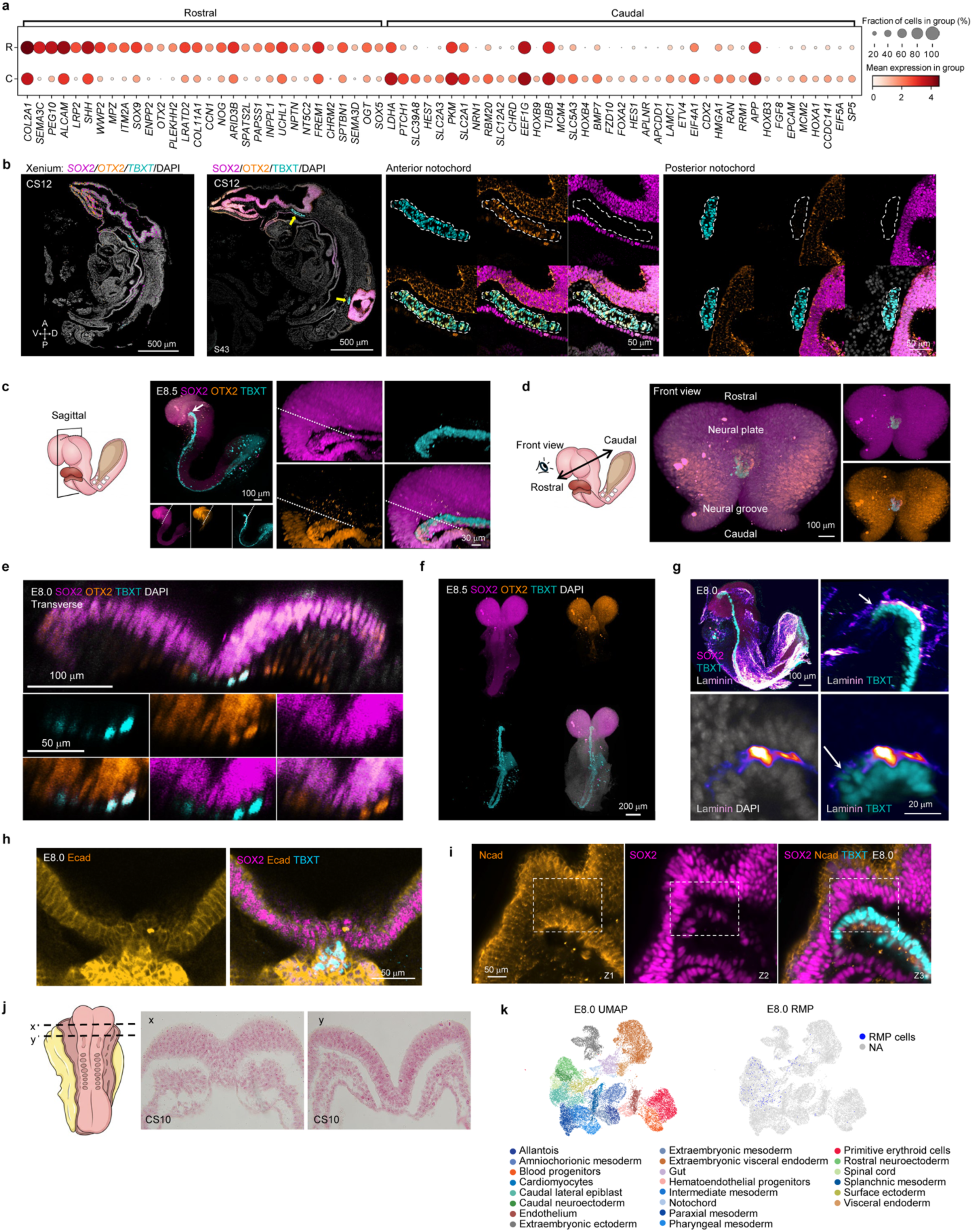
Xenium and immunostaining characterize RMP cells in human and mouse embryos (associated with Figure 4) (a) Dot plot showing the top differentially expressed genes across rostral and caudal notochord populations. (b) Representative Xenium section with *SOX2, TBXT*, and *OTX2* expression is shown at CS12 (left). Immunofluorescence staining for SOX2, TBXT, and OTX2 on the CS12 sections adjacent to the Xenium sections, respectively (middle). Notochord is highlighted by dotted circles. Yellow arrowheads indicate rostral and caudal notochord, respectively. Scale bars as indicated. (**c-d**) Whole-mount immunofluorescence staining of mouse E8.5 embryos for SOX2, OTX2, and TBXT. (left) Schematic of an E8.5 mouse embryo in sagittal and front view, respectively, indicating the neural groove, neural plate, and rostro-caudal axis. Representative sections with SOX2, OTX2, and TBXT staining are shown. Sectioning planes, orientations, and viewing angles are indicated. Scale bars as indicated. (e) Whole-mount immunofluorescence staining of a mouse E8.0 embryo. A representative transverse optical section shows expression of TBXT, SOX2, and OTX2 at the plane indicated. Scale bars as indicated. (f) Whole-mount immunofluorescence staining of a mouse E8.5 embryo for SOX2, OTX2, and TBXT, as shown in (**c**), front view. Scale bar, 200 µm (g) Whole-mount immunofluorescence staining of a mouse E8.0 embryo for SOX2, TBXT, and LAMININ, shown in lateral view and sagittal optical section. Scale bar, 20 µm. (h) Whole-mount immunofluorescence staining of a mouse E8.0 embryo. A representative transverse optical section shows expression of TBXT, SOX2, and E-CADHERIN. Scale bar, 50 µm. (i) Whole-mount immunofluorescence staining of a mouse E8.0 embryo. A representative transverse optical section shows expression of TBXT, SOX2, and N-CADHERIN. The RMP domain is indicated by a dotted box. Z-plane positions (Z) are indicated. Scale bar, 50 µm. (j) Transverse sections of a human CS10 embryo from the Kyoto Collection. The schematic indicates the rostro-caudal levels at which sections were obtained. (k) Left: UMAP embedding of scRNA-seq profiles from E8.0 embryos, colored by annotated cell identity. Right: the same UMAP highlights the distribution of RMP cells in blue dots, showing their localization at the interface of rostral/caudal neuroectoderm and caudal lateral epiblast.

**Supplementary Figure 10.**
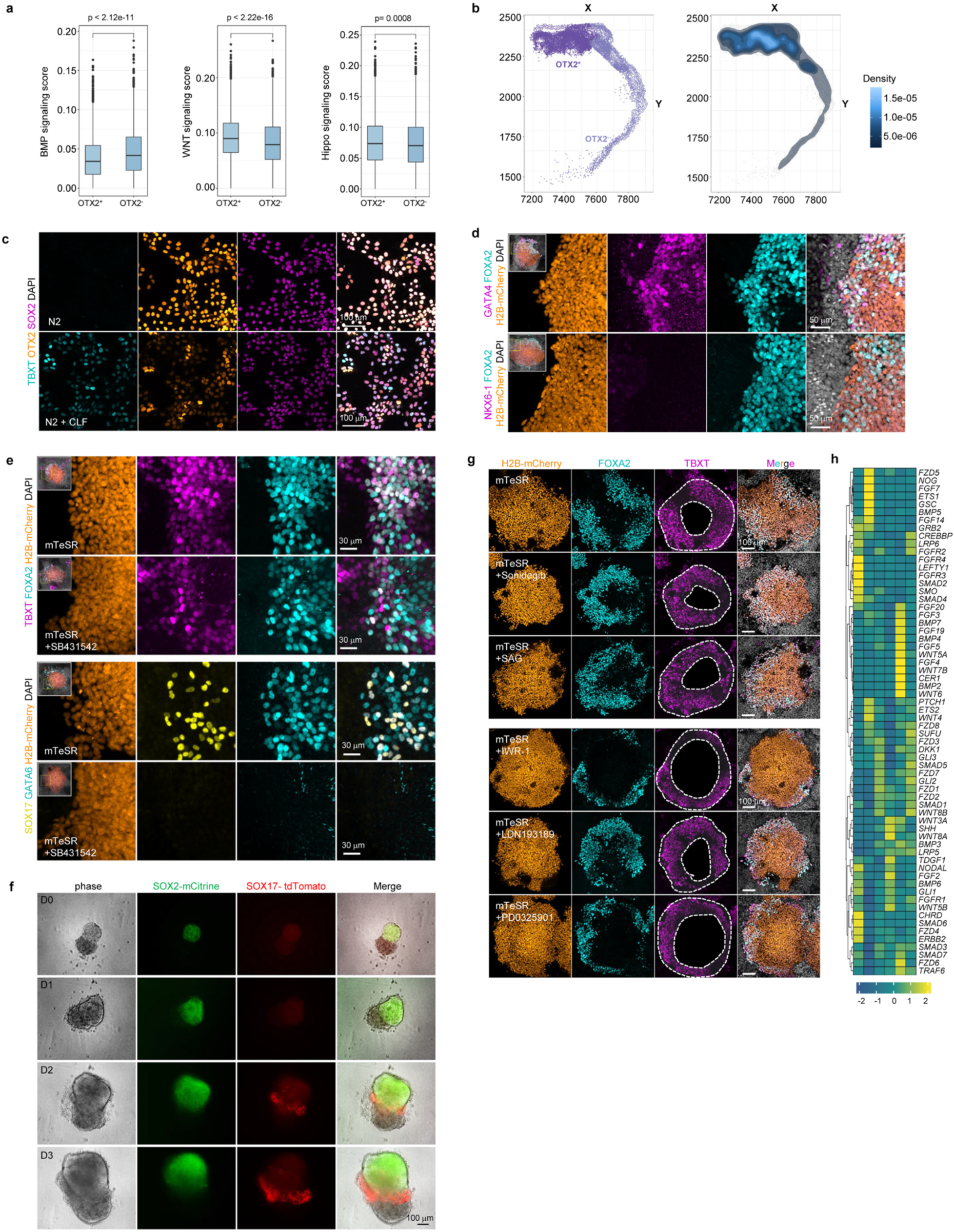
In vitro derivation of RMP-like cells and their inductive capacity (associated with Figure 5) (**a**) BMP, WNT, and Hippo signaling pathway-associated transcriptional scores in OTX2^+^ versus OTX2^-^ neural plate/tube cells from CS9 Visium HD sections shown in (b). OTX2⁺ and OTX2⁻ cells were defined as cells with OTX2 expression above zero and equal to zero, respectively. The pathway-associated transcriptional scores were computed per cell with AUCell using predefined pathway gene sets (Methods; Supplementary Table 5). Boxes show the median and interquartile range; p values, two-sided Wilcoxon rank-sum test. OTX2⁺ cells show higher WNT and Hippo and lower BMP activity than OTX2⁻ cells, a signaling combination matched by the CLF induction medium used to generate RMP-like cells shown in (c). (b) Spatial distribution (left) and local kernel density (right) of OTX2⁺ and OTX2⁻ neural plate/tube cells in the CS9 Visium HD section scored in (a). OTX2⁺ cells occupy a continuous, densely populated rostral domain, indicating a coherent anatomical population rather than scattered detection. The color scale indicates local cell density (c) Induction of RMP-like cells in attachment culture. Day-7 micropatterned hPSCs were treated for 18 h with CLF induction medium (CHIR99021, LDN-193189, and bFGF in N2B27); control cultures were kept in N2B27 without induction. Immunofluorescence staining for SOX2, TBXT, and OTX2 shows that the SOX2⁺TBXT⁺OTX2⁺ state is induced by CLF treatment, whereas control cultures remain TBXT-negative. Scale bar, 100 μm. (d) Immunofluorescence staining of 2D co-cultures of H2B-mCherry-labeled hPSC spheres (right of the dashed box) on RMP-like cell feeders (of the dashed box left) for FOXA2, GATA4, and NKX6-1; after 2 days, Recipient spheres express FOXA2 and GATA4 but remain NKX6-1-negative, indicating that the induction is mesendodermal rather than ventral neural. Scale bar, 50 μm. (e) Immunofluorescence staining of RMP-like cells and hPSC co-culture for TBXT, FOXA2, GATA6, and SOX17 in the absence or presence of the NODAL inhibitor SB431542 shown in zoom-in area. Scale bar, 30 μm. (f) 3D Co-culture of RMP-like cells and hPSCs (RUES2-GLR) for 3 days. SOX2^+^ (mCitrine, green) or SOX17^+^ (tdTomato, red) cells record from co-culture D0 to D3. Scale bar, 100 μm. (g) Co-culture of RMP-like cells and hPSCs in the absence or presence of Sonidegib and SAG (Hedgehog modulation), IWR-1 (WNT inhibition), LDN193189 (BMP inhibition), and PD0325901 (MEK inhibition), stained for FOXA2 and TBXT. The receiver colponies are indicated. Scale bar, 100 μm. (h) Heatmap showing expression of signaling pathway components in RMP-like cell clusters.

**Supplementary Figure 11.**
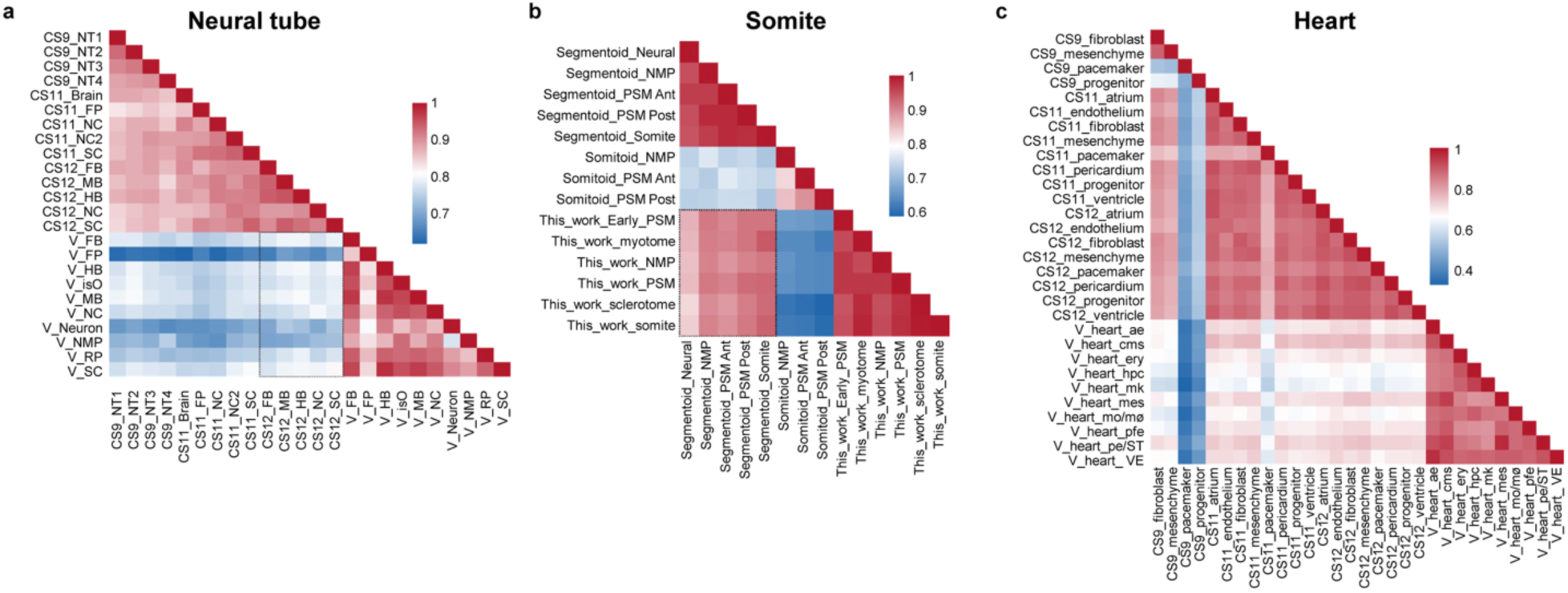
Benchmarking human stem-cell-based embryo models against a spatial transcriptomic atlas. (**a-c**) Heatmaps showing the correlation coefficients between in vivo cell types from CS9, CS11, and CS12 human embryos and the corresponding cell clusters from in vitro neural tube (V), segmentoid, somitoid, and heart models. NP, neural plate; NT, neural tube; NC, neural crest; FP, floor plate; RP, roof plate; Otic, otic placode; Neu prog, neural progenitor; SE, surface ectoderm; LPM, lateral plate mesoderm; FB, forebrain; MB, midbrain; HB, hindbrain; SC, spinal cord; NMP, neuromesodermal progenitors; ST, septum transversum; SplM, splanchnic mesoderm. PSM Ant/Post, anterior/posterior presomitic mesoderm; VE, venous endothelial cells; AE, arterial endothelial cells; HE, haemogenic endothelium; AFE, anterior foregut endoderm; ECs, endothelial cells; CMs, cardiomyocytes; PFE, posterior foregut endoderm; MES, mesenchymal cells; Ery, erythroid cells; Mo/MΦ, monocytes/macrophages; MK, megakaryocytes; HPCs, haematopoietic progenitor cells.

